# New information triggers prospective codes to adapt for flexible navigation

**DOI:** 10.1101/2023.10.31.564814

**Authors:** Stephanie M. Prince, Teema A. Yassine, Navya Katragadda, Tyler C. Roberts, Annabelle C. Singer

## Abstract

Navigating a dynamic world requires rapidly updating choices by integrating past experiences with new information. In hippocampus and prefrontal cortex, neural activity representing future goals is theorized to support planning. However, it remains unknown how prospective goal representations incorporate new, pivotal information. Accordingly, we designed a novel task that precisely introduces new information using virtual reality, and we recorded neural activity as mice flexibly adapted their planned destinations. We found that new information triggered increased hippocampal prospective representations of both possible goals; while in prefrontal cortex, new information caused prospective representations of choices to rapidly shift to the new choice. When mice did not flexibly adapt, prefrontal choice codes failed to switch, despite relatively intact hippocampal goal representations. Prospective code updating depended on the commitment to the initial choice and degree of adaptation needed. Thus, we show how prospective codes update with new information to flexibly adapt ongoing navigational plans.

## INTRODUCTION

The ability to rapidly update our choices in response to new information is essential to navigating a dynamic world. During navigation, animals often hold an internal representation of their world via a cognitive map (Behrens et al., 2018; O’Keefe & Nadel, 1978; Tolman, 1948). When navigation is goal-directed, this internal model is employed to represent not only current state information, but also upcoming choices or possible actions to reach a goal. Current theories propose that these prospective representations of future choices are a neural correlate of planning or deliberation, to simulate consequences of potential actions before they occur (Buckner, 2010; Comrie et al., 2022; Hunt et al., 2021; Lisman & Redish, 2009; Mullally & Maguire, 2014; Pezzulo et al., 2019; Redish, 2016). Such theories imply that these prospective codes facilitate flexible navigation in which animals continuously select from potential choices as events unfold. However, many of the studies testing prospective codes occur in static environments where animals do not need to adapt their behavior within single trials after their initial choices have been made. In dynamic environments where sensory stimuli change frequently, animals must continuously assess new information to inform their decisions and update choices as needed. Thus, it is unclear how prospective codes contribute to flexible navigation that requires continuous assessment of and responses to new information.

Hippocampus, which is essential for spatial navigation, is classically known to encode animals’ current position in an environment, but research has also found that non-local representations occur during active navigation. Prior studies have shown that sweeps of coordinated place cell activity tile the space behind and in front of animals during behavioral hallmarks of deliberation (Johnson & Redish, 2007; Wikenheiser & Redish, 2015). During active behavior, individual neuron activity as well as population-level sequences can represent non-local positions stretching far in front of and behind the animals (Feng et al., 2015; Foster & Wilson, 2007; Gupta et al., 2012; Kapl et al., 2022; Kay et al., 2020; Skaggs et al., 1996; Y. Wang et al., 2014; Wikenheiser & Redish, 2012; Yu & Frank, 2021). One theory is that these sequences and sweeps reflect upcoming choices or might steer behavior toward a specific goal, but the degree to which choices can be predicted from this activity is inconsistent across studies (Wikenheiser & Redish, 2014; Zheng et al., 2021). Another theory is that these non-local position codes are not just representations of animals’ upcoming choices but are instances of hippocampus presenting all possible hypothetical paths. This constant path simulation might allow animals to quickly consider and decide between potential options, a cognitive process that would be especially valuable in dynamic environments where new sensory information changes which potential plans are optimal. Overall, it remains unclear why non-local representations have been observed to sometimes predict upcoming behavior and at other times reflect both options equally.

During goal-directed navigation, animals must not only encode spatial trajectories through their environment, but they must also select and maintain choice information as their trajectories unfold. Prefrontal cortex is thought to represent upcoming choice information and is required for flexible decision-making. In the context of spatial navigation, several studies have found evidence that prefrontal cortex acts in coordination with hippocampus to facilitate decision-making and planning for upcoming choices (Benchenane et al., 2010; den Bakker et al., 2023; Hyman et al., 2005; Jones & Wilson, 2005; O’Neill et al., 2013; Tamura et al., 2017). Prefrontal cortex neural activity in rodents has been observed to co-occur with hippocampal activity specifically when hippocampus encodes non-local spatial representations, suggesting prefrontal cortex may play a role in evaluating these non-local codes as potential choices (Berners-Lee et al., 2021; Hasz & Redish, 2020; Yu & Frank, 2021). Indeed, hippocampal-prefrontal spatial representations in rodents are more coordinated when both regions are representing upcoming choices, and inactivating prefrontal cortex disrupts hippocampal prospective codes and sequences (Guise & Shapiro, 2017; Schmidt et al., 2019; Tang et al., 2021). However, the question remains, what are the roles of hippocampus and prefrontal cortex together in a *dynamic* environment that requires flexible decision-making and reconsideration of potential choices? It is unclear how these hippocampal non-local codes and prefrontal cortex choice representations respond to new information to *accurately* update decisions when plans need to be changed.

To address these open questions, we designed a novel memory-based decision-making task in virtual reality. In this task, we precisely controlled the timing of new information in the environment on a single-trial basis; animals were required to consider two possible paths and flexibly change course mid-trial when new information indicated a switch in the location of the rewarding outcome. We then recorded many units from hippocampus and prefrontal cortex during this behavior in order to evaluate how prospective codes respond to new information that demands flexible navigation. We assessed prospective codes for goal locations and upcoming choices using a Bayesian decoding approach, decoding the animal’s position and evolving choice from neural activity. In hippocampus we found that new, pivotal information causes non-local representations of both possible goal locations to rapidly increase. In prefrontal cortex we found that new, pivotal information causes choice codes to rapidly switch to represent the new choice. We then identified how prospective codes differed when animals failed to adapt; hippocampal position codes remained relatively intact but prefrontal choice codes did not switch to represent the new choice more strongly than the old choice. Finally, we assessed the neural responses as a function of how strongly animals were committed to their choice before new information was presented. We found that both hippocampal and prefrontal cortex prospective codes depended on the animals’ level of commitment to the initial choice and the degree of behavioral adaptation needed. These results reveal how prospective codes support flexible planning and accurate decision-making in the face of new information, and how prospective codes rapidly change to represent both possible options or upcoming choices depending on the degree of behavioral commitment.

## RESULTS

### Animals rapidly update their choices in response to new information in a spatial memory task

To investigate how animals update plans when presented with new, pivotal information, we designed a virtual reality spatial navigation task (**Figure 1A**). Mice (n = 7 animals) were trained to navigate a y-maze using visual cues displayed on the wall (see **Methods** for training paradigm). The first original cue, presented on the walls in the central arm of the track, indicated which arm of the track (left or right) was the rewarded location. On most trials (65%), the original cue then disappeared when mice reached a specific location in the track, and the cue was replaced with uninformative grey checkered walls for the remainder of the trial (delay-only trials). The mice had to maintain the memory of the correct goal arm during the delay period (10.56 ± 0.07 seconds, n = 7172 trials, **Supplementary** Figure 1). On a subset of trials, a second visual cue appeared when mice reached a specific location after a shortened delay period (1.40 ± 0.03 seconds, n = 1840 trials). During the second cue, the visual patterns appeared on the opposite wall from the original cue indicating that the reward location switched from the initial arm, and animals must switch from their initial decision maintained in memory to the opposite choice (switch trials, 25%). The second visual cue was then followed by another delay period before the reward location (6.02 ± 0.08 seconds). After several phases of behavioral training, animals learned to remember and follow the cues across all trial types and performed above 50% accuracy (delay only: 65 ± 2%, switch: 74 ± 2%, p < 0.0001 delay only vs. 50% correct, p < 0.0001 switch vs. 50% correct, **Figure 1B**, **Supplementary** Figure 1B). Performance was slightly worse overall on delay only trials, likely due to the longer delay duration between the original cue and reward compared to update trials. Indeed, behavioral performance varied as a function of delay duration during brief warm-up periods at the start of each session (**Supplementary** Figure 1C). Overall, these results show animals successfully perform both flexible and memory-guided decision-making in this spatial navigation task.

**Figure 1.**
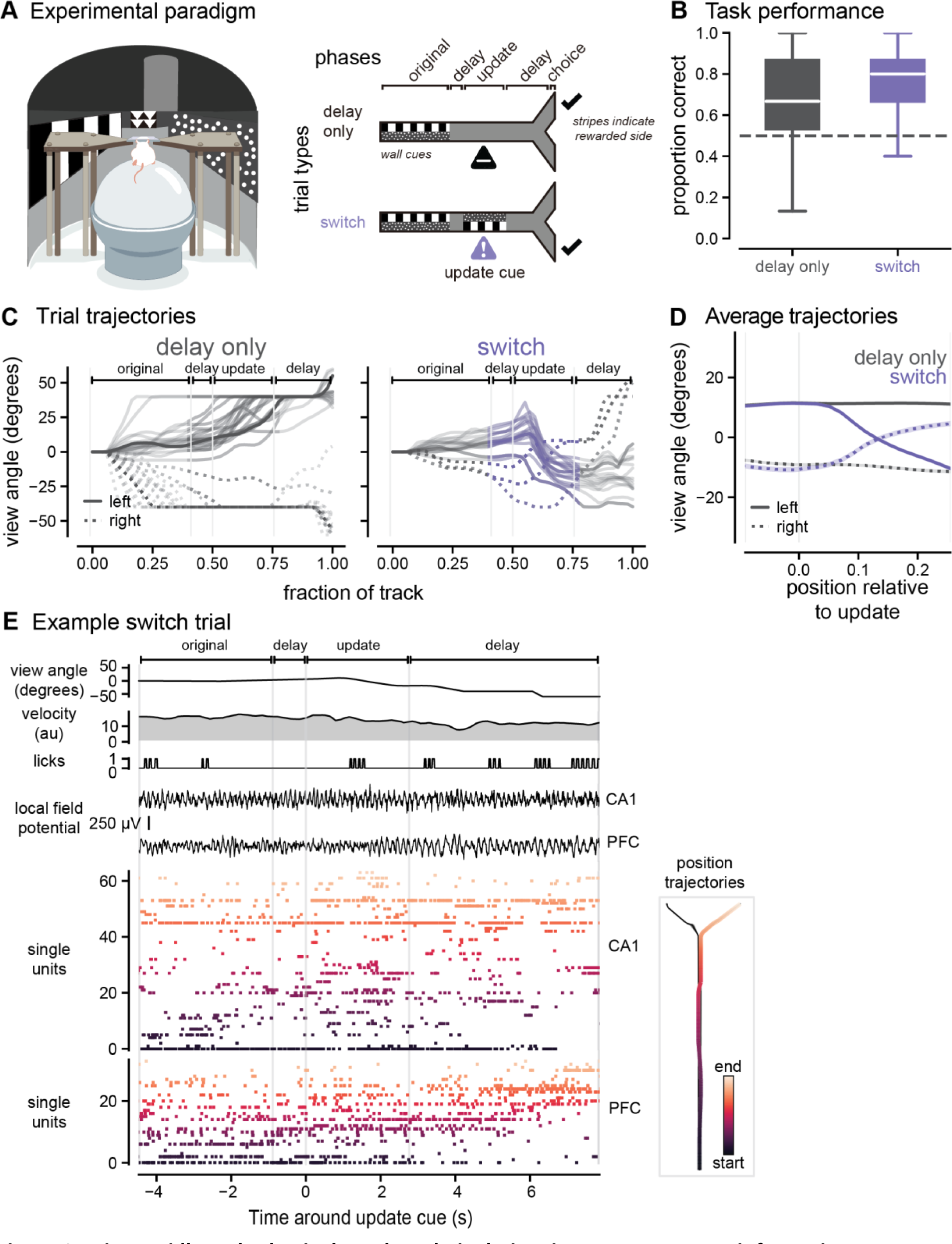
Mice rapidly and selectively update their choices in response to new information. **A.** *Left*, schematic of the virtual reality system. *Right,* update task paradigm with cues displayed along the walls of the track. Delay only trials (black, top, 65% of trials) consisted of an original cue period followed by a delay period during which no cue information was visible. Switch trials (purple, bottom, 25% of trials) consisted of an original cue period, a brief delay period, and then a second cue period during which the wall cues appeared on the opposite side as the original cues. Check marks indicate correct, rewarded side. **B.** Proportion correct on delay only trials (black) and switch trials (purple) for all animals (n = 7 animals). Box plots indicate median and quartiles of distribution of 40-trial bins. (delay only: 0.68 ± 0.00, n = 6252 windows of 40 trials, percentiles = 0.13, 0.53, 0.67, 0.87, 1.00; switch: 0.78 ± 0.01, n = 1002 windows, percentiles = 0.27, 0.67, 0.80, 0.87, 1.00). Linear mixed effects model (LME), see Table 2 for statistical details. **C.** Example behavioral trajectories for individual trials of each trial type. View angle (heading direction) across locations in the track for example correct left (solid lines) and right (dashed lines) trials. Task phases indicated with brackets and light grey vertical lines. Heading direction is set at 0 at trial onset. On update trials, the delay period preceding the update location and the update cue location are shown in purple. **D.** Average view angle trajectories across all delay (grey) and switch (purple) trials. Dashed lines indicate initial right trials, solid lines indicate initial left trials, mean + SEM. **E.** An example trial and simultaneous electrophysiological recordings from hippocampal CA1 and medial prefrontal cortex. *Left*, local field potential from the channel with the peak sharp-wave ripple power in hippocampal CA1. Single units, including putative pyramidal cells and interneurons, sorted by spatial tuning, darker colors indicate place fields earlier in the track, lighter orange colors indicate place fields at the end of the track. Grey vertical lines indicate timepoints in the task, data shown from the moment animals movement in the virtual environment is unrestricted (3 seconds after trial start) to the moment animals reach the end of the environment and have made a choice. *Right*, plots of the 2-dimensional position trajectory for the same example trial. Darker orange colors indicate earlier in the spatial trajectory and lighter colors indicate the end. Other spatial trajectories for the same behavioral session shown in black in the background. Note there is minimal lateral displacement until the goal arms due to the restrictions of the virtual environment ns, not significant, *p < 0.05, **p < 0.01, ***p < 0.001, ****p < 0.0001. All values reported as mean ± SEM, percentiles represent [minimum, 25^th^, median, 75^th^, maximum].

Given our central question about how prospective codes respond to flexible navigation demands, we were specifically interested in the behavior around the update cue on the switch trials, when animals had to update their behavioral trajectories in response to new information. The animals’ heading direction (view angle, see **Methods**) slowly diverged on average as animals approached the ends of the track (**Figure 1C**). We found that on switch trials around the update cue onset, the animals’ heading direction shifted from one direction to the opposite in response to the cue indicating the reward location had changed (**Figure 1D**, see **Supplementary** Figure 1D for other behavioral metrics). These behavioral data show that animals rapidly switch decisions in response to new information.

### Enhanced non-local codes of both goal locations in hippocampus in response to new, pivotal information

To determine how prospective codes change with new information in a dynamic environment, we recorded single unit and local field potential activity in dorsal hippocampal CA1 and medial prefrontal cortex during behavior (n = 3892 units in hippocampus, n = 2557 units in medial prefrontal cortex, **Figure *1*E**, **Supplementary** Figure 2A-B, **Table 1**). As expected from prior literature, we found that neurons in both hippocampus and prefrontal cortex had spatial tuning curves that tiled the virtual environment (**Supplementary** Figure 2C**, Supplementary** Figure 4A). To determine if prospective codes of position represent possible paths or planned paths in response to new information, we estimated population-level representations of position using memoryless Bayesian decoding of hippocampal and prefrontal activity (see **Methods**). We first confirmed that hippocampal and prefrontal activity could reliably decode the animals’ current location in the environment; decoded positions from neural activity represented the animals’ true position across locations in the environment (**Supplementary** Figure 2D**, Supplementary** Figure 4B).

We used this decoding approach to understand how neural representations change when animals receive new information that directs them away from a previous goal destination to a new goal location. To do so, we focused specifically on neural activity around the update cue onset. We expected that neural activity in hippocampus would predominantly represent animals’ current position, but there would be some remote prospective position coding (Jezek et al., 2011; Kapl et al., 2022; Kay et al., 2020; Yu & Frank, 2021). We hypothesized that the amount of non-local representation would remain constant, and both goal locations would be represented equally in hippocampus while animals were in the central arm of the track, even when new information was presented. Before the update cue onset and on delay-only trials, decoded positions predominantly represented the animals’ current position (**Figure *2***). In contrast, we were surprised to observe that shortly after the update cue was presented on switch trials, the decoding output rapidly jumped from predominantly representing the animals’ current location, to representing non-local locations within both the new and initial goal arms (**Figure 2A-B, Supplementary** Figure 2E). We quantified this elevated non-local representation and found that the overall probability of goal locations being decoded, the posterior probability integrated over each goal arm, was elevated on switch trials compared to delay only trials (p < 0.0001, initial, switch vs. initial, delay only, p < 0.0001, new, switch vs. new, delay only, in order to assess statistical significance while appropriately accounting for repeated measures within animals and within sessions, we used a linear mixed effects model (LME) approach, see **Methods** and **Table 2** for additional statistical details, **Figure 2C-D, Supplementary** Figure 3). This *elevation* of non-local coding in response to new, crucial information was contrary to our expectations, because we predicted the amount of non-local coding would remain relatively stable throughout the trial similar to what we observed during delay only trials. We then wondered if both goals were equally represented, consistent with a role for prospective codes representing possible paths. We found both the new correct goal location and the initial correct goal location increased similar amounts on average. When we compared differences between the amount of initial goal location and new goal location representations on a trial-by-trial basis, the differences were not significantly different from zero and were not significantly different between switch and delay only trials (p = 0.7806 switch vs. delay only, **Figure 2D**, see **Table 2** for additional statistical details). Overall, these results show that when new, crucial information is presented that requires animals to update their goal destinations, hippocampus suppresses local position coding and increases non-local coding of *both* potential goal locations.

**Figure 2.**
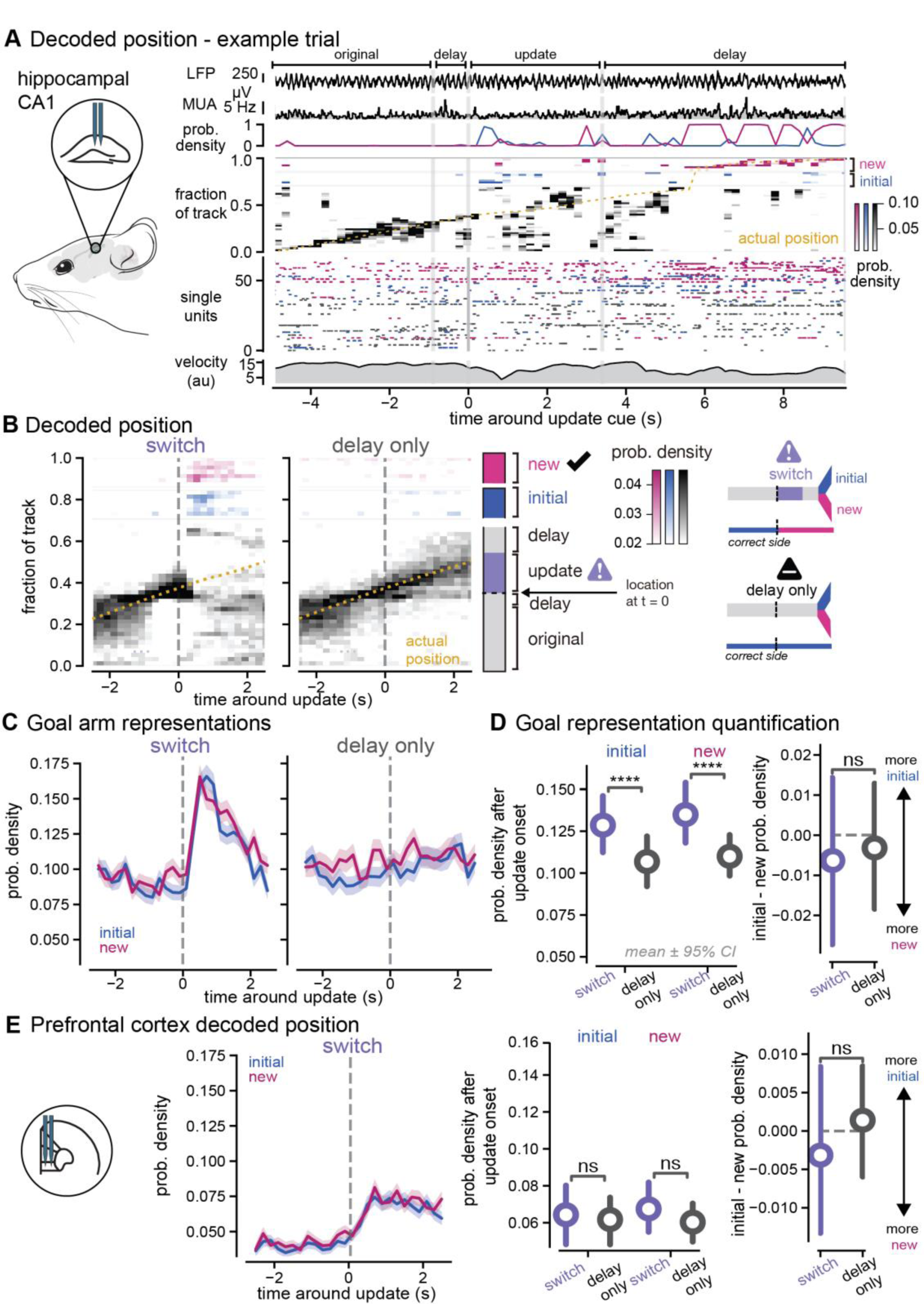
Non-local representations for both initial and new goals increase in hippocampus in response to new information **A.** *Left*, schematic of hippocampal CA1 recording location. *Right*, decoding of position from hippocampal spiking activity on an example update trial. *First row*, local field potential from hippocampal CA1. *Second row*, multi-unit activity from recorded single units. *Third row*, probability density of goal arms being decoded for each timepoint of the trial. *Fourth row*, heatmap indicating the posterior probability density function of spatial locations decoded from neural activity in 200 ms time bins. *Fifth row*, single unit activity used as inputs into the Bayesian decoder. Units sorted by place field peak and color coded with the location of the largest place field (in the central arm (black), initial arm (blue), or new arm (pink)). *Sixth row*, summed rotational and translational velocities from the spherical treadmill. Grey vertical lines indicate timepoints in the task. **B.** *Left,* Decoding output (posterior probability density) before and after the update cue is presented. Average across all recording sessions for switch trials (*center*) and delay only trials (*right*). Heatmap indicates which positions are most strongly decoded by the spiking activity of all hippocampal neurons. Pink and blue indicate decoding of positions within the new and initial goal arms (including the entire goal arms after the choice point), which occurs during the update cue even though the animals are in the center arm away from the goal locations. Animals’ actual position (yellow dashed line) shown over the same time window. Data from both left and right trial types were combined. Two-dimensional position in the virtual track was converted to a 1D position (see ***Methods***) to disambiguate the new and initial arms of the environment. *Middle*, schematic of locations of the task as shown on y-axis of left panels. *Right*, schematic of trial types illustrating initial (blue) and new (pink) goal locations and the update cue onset (purple). On switch trials the correct goal switches from the initial to new arm, while on delay trials the correct goal arm is always the initial arm. **C.** Decoding (integrated probability densities) of all locations within the new (pink) and initial (blue) goal arms around the update cue on switch (center) and delay only (right) trials. Dark pink and blue lines indicate bins significantly different from baseline. Mean ± SEM across all trials shown. **D.** *Left*, probability density decoding of the initial (left) and new (right) goal locations in the first 1.5 seconds after the update cue onset on switch (purple) and delay only (black) trials. (initial, switch: 0.13 ± 0.00, n = 1295 trials, percentiles = 0.00, 0.03, 0.10, 0.19, 0.87; new, switch: 0.14 ± 0.00, n = 1295 trials, percentiles = 0.00, 0.04, 0.10, 0.21, 0.97; initial, delay only: 0.10 ± 0.00, n = 926 trials, percentiles = 0.00, 0.02, 0.07, 0.15, 0.88; new, delay only: 0.11 ± 0.00, n = 926 trials, percentiles = 0.00, 0.02, 0.08, 0.16, 1.00). Colored circle and line indicate mean ± 95% CI computed with n = 1000 bootstrap samples. *Right,* difference on single trials between initial and new probability densities after the update cue. (switch: -0.01 ± 0.01, n = 1295 trials, percentiles = -0.95, -0.11, 0.00, 0.10, 0.87; delay only: -0.01 ± 0.01, n = 926 trials, percentiles = -0.99, -0.08, -0.00, 0.07, 0.88). Linear mixed effects model (LME) and ANOVA Tukey post-hoc pairwise test, see Table 2 for statistical details. **E.** As in **C** and **D** for prefrontal cortex position decoding on switch trials. Probability density decoding of initial and new goal locations (initial, switch: 0.07 ± 0.00, n = 1109 trials, percentiles = 0.00, 0.00, 0.03, 0.09, 0.80; new, switch: 0.07 ± 0.00, n = 1109 trials, percentiles = 0.00, 0.00, 0.03, 0.09, 0.88; initial, delay only: 0.06 ± 0.00, n = 798 trials, percentiles = 0.00, 0.00, 0.03, 0.08, 0.72; new, delay only: 0.06 ± 0.00, n = 798 trials, percentiles = 0.00, 0.00, 0.03, 0.09, 0.68). Difference on single trials between initial and new probability densities (delay only: -0.00 ± 0.00, n = 798 trials, percentiles = -0.60, -0.02, 0.00, 0.02, 0.71; switch: -0.00 ± 0.00, n = 1109 trials, percentiles = -0.83, -0.03, -0.00, 0.02, 0.76). Linear mixed effects model (LME) and ANOVA Tukey post-hoc, see Table 2 for statistical details. ns, not significant, *p < 0.05, **p < 0.01, ***p < 0.001, ****p < 0.0001. All values reported as mean ± SEM, percentiles represent [minimum, 25^th^, median, 75^th^, maximum].

To confirm this elevation of non-local coding was not due to a complete loss of accurate decoding, we compared the representation of the goal arms to a uniform decoding output, that is the theoretical posterior probability value if all locations in the track were equally represented. We found that the elevated probability in hippocampus was 15% and 19% greater than uniform decoding for initial and new goal locations. Furthermore, this increased non-local activity was not due to overall poor decoding of that part of the environment. On delay only trials, the neural activity continued to predominantly represent the animals’ current location in the central arm of the track over the same time interval (85% greater than uniform decoding on delay only trials, **Figure *2*B**). We trained our encoding model with delay only trials, so that the spatial tuning curves were not influenced by the neural activity that occurred on the update trials and we could accurately assess how neural activity varied on update trials compared to delay only trials. However, we wondered if these results showing increased non-local decoding after the update were due to the fact that our model was trained on delay only trials. To control for this possibility, we also ran the analysis using all trials in the encoding model and found that while there was more accurate representation of the animals’ current location during the update cue, the non-local coding results remained similar (data not shown). Because the update trials are interleaved with the delay only trials, it is unlikely that these differences are due to spatial remapping, single unit instability, or behavioral disengagement.

In prefrontal cortex, we expected that decoding position from neural activity would also reveal representation of primarily the animals’ current position with some weaker representation of the remote goal arms. We hypothesized that in the update task, prospective codes for goal locations in prefrontal cortex would switch from the initial goal location to a new one when animals switched their behavioral trajectory. We found that the overall decoding output was much less spatially specific in medial prefrontal cortex than hippocampus. Furthermore, neural activity represented the animals’ current position more than distal locations (**Supplementary** Figure 4A-C). Non-local representations in medial prefrontal cortex were not significantly different across trial types and both goal locations were equally represented until the animals approached the goal arm (p = 0.3496, initial, switch vs. initial, delay only, p = 0.3148, new, switch vs. new, delay only, p = 0.9111, switch vs. delay only, **Figure 2E, Supplementary** Figure 4D). In summary, these results show that when new information is presented that requires an update in goal destination, unlike hippocampus, prefrontal cortex spatial representations continue to predominantly represent the animals’ current position and do not seem to diverge in representing one goal location over the other until the animals approach the choice point.

### Theta phase segregation of non-local and local position codes

Prior studies have shown that local and non-local codes are segregated into different phases of theta oscillations, providing temporal organization and segregation of these different spatial representations. Given the increase of non-local coding we found in hippocampus, we wondered if current and future location codes were segregated by theta phase, as in prior work in static environments without updates (Feng et al., 2015; O’Keefe & Recce, 1993; Skaggs et al., 1996; Wikenheiser & Redish, 2015). We hypothesized that codes for prospective locations would increase at a specific phase of theta, separate from the phase where local codes dominate, when animals were presented with new information that required them to reconsider potential choices. Using our population-level metrics, we identified the theta phase of each decoded time bin and the corresponding location in the environment (initial goal arm, new goal arm, or central arm). In line with previous work, we found that codes for prospective goal locations (non-local) were represented significantly more on the opposite theta phase as current location representations (local) (p = 0.0275, -π to 0, initial vs. central arm, p = 0.0106, -π to 0 new vs. central arm, p < 0.0001, 0 to π, initial vs. central arm, p < 0.0001, 0 to π, new vs. central arm, **Figure 3A-B**). Specifically, non-local representations were highest in the first half of the theta cycle, while local representations were highest during the second half of the theta cycle. When comparing before and after the update cue onset, we found that there was a significant increase in the amount of goal location (non-local) decoding, but the local vs. non-local phase preferences were maintained (p < 0.0001, pre- vs. post-update initial arm, p < 0.0001 pre- vs. post-update new goal, **Figure 3B-C**). Similar to our behavioral-timescale findings, non-local codes at the timescale of individual theta cycles were also significantly larger on switch trials compared delay-only trials (p < 0.0001, initial arm, switch vs. delay only, p < 0.0001 new arm, switch vs. delay only **Supplementary** Figure 5). However, in contrast to our longer timescale results, we observed slight differences in initial and new goal codes after the update cue. The new goal location was more strongly theta modulated compared to the old goal location (p = 0.0008, new arm -π to 0 vs 0 to π, p = 0.6496, initial arm arm -π to 0 vs 0 to π, effects were similar when breaking down by quarters, **Figure *3*C**). Because different locations can be represented on different theta cycles (theta cycle skipping), we also hypothesized that such theta cycle skipping would increase when animals were presented with new information. However, the rates of theta cycle skipping of individual neurons were similar across switch and delay only trials (data not shown). Overall, these results show that even with large increases in non-local coding and decreased local coding at longer timescales, non-local goal representations remain segregated from local codes during theta oscillations. While initial and new goals are representated at similar levels in hippocampus at both a behavioral and theta-timescale, stronger theta modulation of new goal codes in particular after the update cue onset suggests more organized spike timing of new goals in response to new, pivotal information.

**Figure 3.**
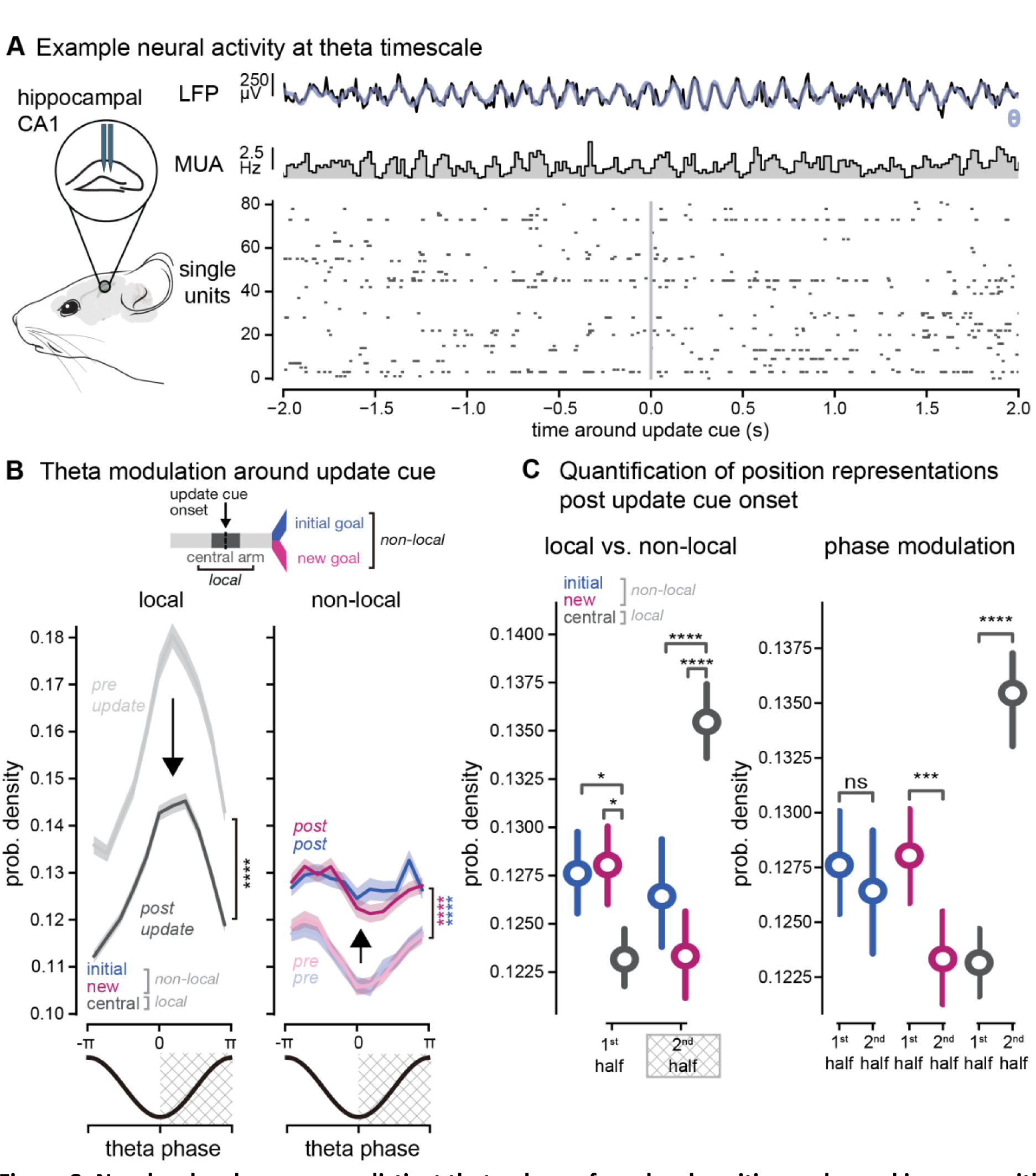
Non-local codes occur on distinct theta phases from local position codes and increase with new information **A.** Example local field potential and single unit activity at the theta timescale. *First row*, example theta oscillation detected in hippocampal CA1 from the local field potential. Low pass filtered signal in black, theta band signal in blue. *Middle row,* multi-unit activity from recorded single units. *Bottom row*, single unit spiking activity. **B.** *Top*, schematic of central arm (local location at time analyzed) and initial and new goal arm (non-local) bins used to quantify local and non-local representations by theta phase. *Bottom*, average decoded central arm (black), initial arm (blue), and new arm (pink) probability densities as a function of theta phase for local (left) and non-local codes (right). Pre-update cue (-1.5 to 0 seconds) shown in lighter shades, post cue (0 to 1.5 seconds) shown in darker shades. Data shown from 1.5 second time period before and after the update cue onset, time window based on previous longer timescale elevation in non-local hippocampal codes from Figure 2. Percentiles for first half of theta cycle, pre-post comparisons similar for second half: (pre, initial: 0.11 ± 0.00, n = 1295 trials, percentiles = 0.01, 0.09, 0.11, 0.13, 0.31; post, initial: 0.13 ± 0.00, n = 1295 trials, percentiles = 0.02, 0.10, 0.12, 0.15, 0.34; pre, new: 0.11 ± 0.00, n = 1295 trials, percentiles = 0.01, 0.09, 0.11, 0.13, 0.30; post, new: 0.13 ± 0.00, n = 1295 trials, percentiles = 0.02, 0.11, 0.12, 0.15, 0.38; pre, central: 0.15 ± 0.00, n = 1295 trials, percentiles = 0.03, 0.12, 0.13, 0.17, 0.37; post, central: 0.12 ± 0.00, n = 1295 trials, percentiles = 0.01, 0.11, 0.12, 0.14, 0.30). Mean ± SEM across all trials shown. Linear mixed effects model (LME) and ANOVA Tukey post-hoc, see Table 2 for statistical details. **C.** *Left,* probability density of the central (local), new (non-local), and initial (non-local) arm in the first and second halves of the theta cycle (1^st^ half indicates -π to 0 and 2^nd^ half indicates 0 to π) after the update cue onset on switch trials. *Right,* as on left with statistical comparisons between phases. Initial, -π to 0 : 0.13 ± 0.00, n = 1295 trials, percentiles = 0.02, 0.10, 0.12, 0.15, 0.34; initial, 0 to π: 0.13 ± 0.00, n = 1295 trials, percentiles = 0.02, 0.10, 0.12, 0.14, 0.45; new, -π to 0: 0.13 ± 0.00, n = 1295 trials, percentiles = 0.02, 0.11, 0.12, 0.15, 0.38; new, 0 to π: 0.12 ± 0.00, n = 1295 trials, percentiles = 0.01, 0.10, 0.12, 0.14, 0.43; central, -π to 0: 0.12 ± 0.00, n = 1295 trials, percentiles = 0.01, 0.11, 0.12, 0.14, 0.30; central, 0 to π: 0.14 ± 0.00, n = 1295 trials, percentiles = 0.02, 0.12, 0.13, 0.15, 0.45. Linear mixed effects model (LME) and ANOVA Tukey post-hoc, see Table 2 for statistical details. ns, not significant, *p < 0.05, **p < 0.01, ***p < 0.001, ****p < 0.0001. All values reported as mean ± SEM, percentiles represent [minimum, 25^th^, median, 75^th^, maximum].

### Rapid switch from old to new choice estimates in prefrontal cortex when new information is presented

While we were surprised that neural activity in prefrontal cortex did not represent the future goal location more than the alternative goal, this may be explained by the overall minimal non-local coding we observed in prefrontal cortex around the update. Because medial prefrontal cortex is known to encode neural correlates of choice in decision-making tasks, we pursued a different approach to explicitly test how the update cue altered choice codes in prefrontal cortex. We hypothesized that in medial prefrontal cortex the choice representation would be stronger for the initial choice preceding the update cue and would then shift from the initial choice to the new choice after the new information was presented. To test this hypothesis, we took advantage of the continuous development of choice that occurs as animals run down the virtual reality environment and is reflected in the animals’ changing velocity and heading direction over the course of the track (Tseng et al., 2022). To obtain a continuous estimate of relative choice commitment throughout the trial, we trained a long short-term memory (LSTM) neural network to predict the mouse’s ultimate choice from only the behavioral readouts, specifically velocity, heading direction, and forward position trajectories throughout the trial (**Figure 4A**). At each timepoint in the trial, all previous trial time points were used to estimate the animals’ final choice. The choice estimate varied throughout the course of a trial, starting at chance-level accuracy at the start of the trial on average, and ending with a highly accurate, or -0.17 ± 0.02 log likelihood on average, prediction of the animals’ final choice, where 0 is perfect and -1 is a chance level prediction accuracy (**Figure 4A**). We then used this choice measure to obtain a population level estimate of the animals’ choice from medial prefrontal cortex using a Bayesian decoder, similar to our previous decoding of position representation (**Supplementary** Figure 6A-B).

**Figure 4.**
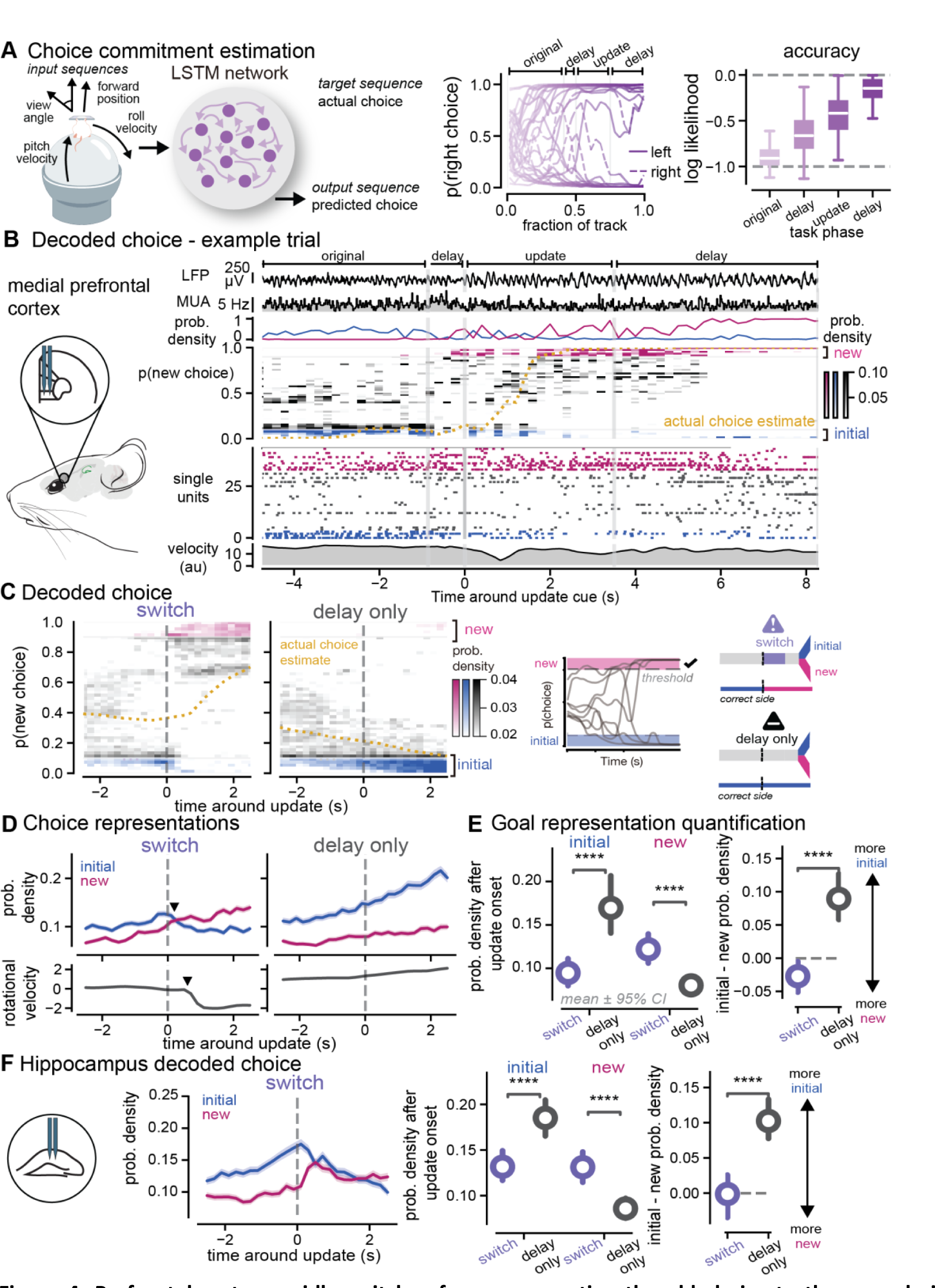
Prefrontal cortex rapidly switches from representing the old choice to the new choice in response to new information. **A.** *Left,* Schematic of long short-term memory (LSTM) neural network for predicting the animals’ choice from the behavioral trajectories. *Center,* example trials of the LSTM output for trials with left (solid) and right (dashed) reported choices. *Right,* session-averaged overall accuracy of the LSTM choice output throughout the virtual environment. (original cue: -0.98 ± 0.05, n = 66 session averages, percentiles = -2.62, -0.97, -0.91, -0.82, -0.60; delay cue: -0.63 ± 0.03, n = 66 session averages, percentiles = -1.13, -0.79, -0.66, -0.50, -0.07; update cue: -0.43 ± 0.03, n = 66 session averages, percentiles = -1.07, -0.58, -0.41, -0.29, -0.01; delay2 cue: -0.17 ± 0.02, n = 66 session averages, percentiles = -0.72, -0.23, -0.14, -0.06, -0.00). **B.** *Left*, schematic of medial prefrontal cortex recordings. *Right*, decoding of choice from prefrontal spiking activity on an example update trial. *First row*, local field potential from prefrontal cortex. *Second row*, multi-unit activity from recorded single units. *Third row*, probability density of new (pink) and initial (blue) choice being decoded for each timepoint of the trial. *Fourth row*, heatmap indicating the probability density function of choices decoded from neural activity in 200 ms time bins. *Fifth row*, single unit activity used as inputs into the Bayesian decoder. Units sorted by tuning curve peak and color coded with the choice preference of no choice preference (black), initial choice (blue), or new choice (pink). *Sixth row*, summed rotational and translational velocities from the spherical treadmill. Grey vertical lines indicate timepoints in the task. **C.** *Left*, decoding output (posterior probability density) of choice before and after the update cue, averaged across all recording sessions. Heatmap indicates stronger decoding of that choice by the spiking activity of all prefrontal neurons. Animals’ actual averaged choice estimate (yellow dashed line) shown over the same time window. Data from both left and right trial types were combined. *Middle*, schematic of choice estimate. The threshold indicates the highest 10% of all choice estimate values for either choice to select fo strong choice decoding of the initial and new choices. *Right*, schematic of trial types illustrating initial (blue) and new (pink) choices around the update cue onset (purple). On switch trials the correct goal switches from the initial to new arm, while on delay trials the correct goal arm is always the initial arm. **D.** *Top*, probability densities of the new (pink) and initial (blue) choice estimates around the update cue on switch (left) and delay only (right) trials. Arrow indicates moment when new cue probability density is greater than the initial cue. *Bottom*, rotational velocity of the spherical treadmill around the update cue. Arrow indicates the first time after the update cue that the change in velocity slope is negative, signifying a change in direction. Mean ± SEM across all trials shown. **E.** *Left*, Probability density of decoding initial (left) and new (right) choices after the update cue on switch (purple) and delay only (black) trials. Colored circle and line indicate mean ± 95% CI computed with n = 1000 bootstrap samples. (initial, delay only: 0.17 ± 0.01, n = 798 trials, percentiles = 0.00, 0.05, 0.11, 0.23, 0.97; initial, switch: 0.10 ± 0.00, n = 1109 trials, percentiles = 0.00, 0.03, 0.07, 0.13, 0.85; new, delay only: 0.09 ± 0.00, n = 798 trials, percentiles = 0.00, 0.02, 0.06, 0.12, 0.71; new, switch: 0.12 ± 0.00, n = 1109 trials, percentiles = 0.00, 0.04, 0.09, 0.16, 0.69). *Right*, Difference between initial and new probability densities after the update cue. (delay only: 0.08 ± 0.01, n = 798 trials, percentiles = -0.71, -0.02, 0.04, 0.16, 0.97; switch: -0.02 ± 0.01, n = 1109 trials, percentiles = -0.66, -0.09, -0.01, 0.05, 0.83). Linear mixed effects model (LME) and ANOVA Tukey posthoc, see Table 2 for statistical details. **F.** As in **D** and **E** for hippocampal choice decoding on switch trials (initial, delay only: 0.19 ± 0.01, n = 885 trials, percentiles = 0.00, 0.07, 0.14, 0.25, 0.94; initial, switch: 0.14 ± 0.00, n = 1238 trials, percentiles = 0.00, 0.05, 0.11, 0.19, 0.80; new, delay only: 0.09 ± 0.00, n = 895 trials, percentiles = 0.00, 0.03, 0.07, 0.12, 0.92; new, switch: 0.13 ± 0.00, n = 1262 trials, percentiles = 0.00, 0.05, 0.10, 0.17, 0.91). Difference between initial and new probabilitiy densities after the update cue (delay only: 0.10 ± 0.01, n = 854 trials, percentiles = -0.86, -0.02, 0.06, 0.19, 0.94; switch: 0.01 ± 0.01, n = 1205 trials, percentiles = -0.91, -0.08, 0.01, 0.10, 0.77). Linear mixed effect model (LME) and ANOVA Tukey posthoc, see Table 2 for statistical details. ns, not significant, *p < 0.05, **p < 0.01, ***p < 0.001, ****p < 0.0001. All values reported as mean ± SEM, percentiles represent [minimum, 25^th^, median, 75^th^, maximum].

Using this approach to decode choice from prefrontal activity, we tested our hypothesis that new information causes choice representations in prefrontal cortex to rapidly switch from the old to the new choice. We observed that the choice estimates of the initial and new choice began to diverge preceding the update and then rapidly flipped after the reward location changed (p < 0.0001 initial switch vs. initial, delay only, p < 0.0001 new, switch vs. new, delay only **Figure 4B-D, Supplementary** Figure 6C**, Supplementary** Figure 7). Interestingly, on update trials the neural activity rapidly switched from high decoding of the initial choice to high decoding of the new choice, with minimal time spent showing no strong choice estimate (**Figure 4C**). Furthermore, the difference between initial and new choice decoding was significantly different between switch and delay trial types (p < 0.0001 initial – new for switch vs. delay only **Figure 4E**). On switch trials, after the update cue the new choice decoding was larger than initial choice decoding on average (resulting in a difference that was below zero). Meanwhile on delay only trials, the initial choice decoding was larger than the new choice during the delay where no new information was presented. We compared the choice representations to a uniform decoding output, that is the theoretical posterior probability value if all choice values were equally represented. We found that the elevated probability in prefrontal cortex was 19% greater than uniform decoding for new choices after the update. On average, the change in neural activity choice decoding preceded the movement change (**Figure 4D**). Examining when animals began changing head direction (rotational velocity began decelerating away from the initial choice) on a single trial basis, we found initial choice codes decreased and new choice codes increased before the motor response (**Supplementary** Figure 6D). Subsequently, the new choice decoding output overtook the initial choice decoding at the same time as the motor response (**Supplementary** Figure 6D). Thus, these changes in neural decoding likely precede motor responses and these effects are not likely due to motor response execution. Overall, these results show that when animals are presented with new information that required a switch from one choice to another, the new choice is quickly represented more than the initial choice, compared to trials that do not require behavioral flexibility. In contrast to prospective codes for positions in the hippocampus that represented possible paths, these results are consistent with hypothesized roles for prospective codes in guiding behavior.

While the hippocampus is traditionally associated with position more than choice representations, prior work has shown that choice can be decoded from hippocampal activity during decision-making tasks. We wondered how new information would affect decoded choice estimate representations in hippocampus (**Supplementary** Figure 8A-B). Given our previous results showing equal hippocampal position representations of both goals, we hypothesized that both choices would be represented equally in hippocampus after the update. We found that initial choice representations in hippocampus were higher than the alternative choice preceding the update, and then decreased after the update cue was presented and the new choice representation increased (**Figure 4F**, **Supplementary** Figure 8C-D). However, while the decoded activity followed a similar pattern of switching from the initial to the new choice as in prefrontal cortex, this change evolved over a longer period of time in hippocampus. There was an extended period in which both choices were equally represented in hippocampus, but the choice information had already flipped in medial prefrontal cortex. When quantifying the difference between the amount of initial and new choice representations in hippocampus after the update cue was presented, both choices were represented equally and the difference on a trial-by-trial basis was not significantly different from zero (p = 0.5625, initial – new vs. zero, **Figure 4F**) Overall, these findings show that new crucial information that requires animals to update their choices results in a switch from old to new choice representations in both hippocampus and prefrontal cortex. However, the representation of both choices converges to similar levels in hippocampus for an extended period of time, while the representation of the new choice more rapidly overcomes the old choice in prefrontal cortex.

### Goal codes for updating behaviors are not due simply to new visual information

One of the key concerns we had in the interpretation of our results was whether these differences in goal coding were due simply to the visual change that occurred when the update cue turned on and new information was presented. To control for this possibility, we included an additional set of trials that had the same structure as the switch trials, but the second visual cue was on the same side of the track as initially shown, providing additional evidence that the initial reward location was still correct (stay trials, 10%, **Figure 5A-B**). Thus, stay trials included a similar visual cue as on switch trials but no behavioral change was needed. Examining hippocampal position coding, we found that while there was a transient increase in non-local information on stay trials, the amount of non-local representations on stay trials was not significantly different from delay-only trials (p = 0.9927, initial, stay vs. delay only; p = 0.6267, new, stay vs. delay only). Non-local codes were in fact significantly lower on stay trials compared to switch trials (p < 0.0001 initial, stay vs. switch, p = 0.0274 new, stay vs. switch **Figure 5C-D**). Next, we analyzed prefrontal choice coding on stay trials. We found that new choice coding on stay trials was also at similar levels to delay only trials, and significantly lower than switch trials (p = 0.7914, stay vs. delay only, p = 0.0072, stay vs. switch, **Figure 5E-F**). The decoded initial choice estimate remained relatively constant on stay trials, in contrast to switch trials, though also significantly lower than on delay trials (p = 0.0032, stay vs. delay only, p < 0.0001 stay vs. switch). Furthermore, the difference between initial and new choice decoding was significantly different between trial types (p < 0.0001 initial – new for switch vs. stay, p = 0.0033 initial – new for stay vs. delay only). Overall, these results show that the observed changes in goal codes in hippocampus and prefrontal cortex are not due only to the presence of *any* additional visual information, but specifically to new, *pivotal* information that requires animals to update their behavior.

**Figure 5.**
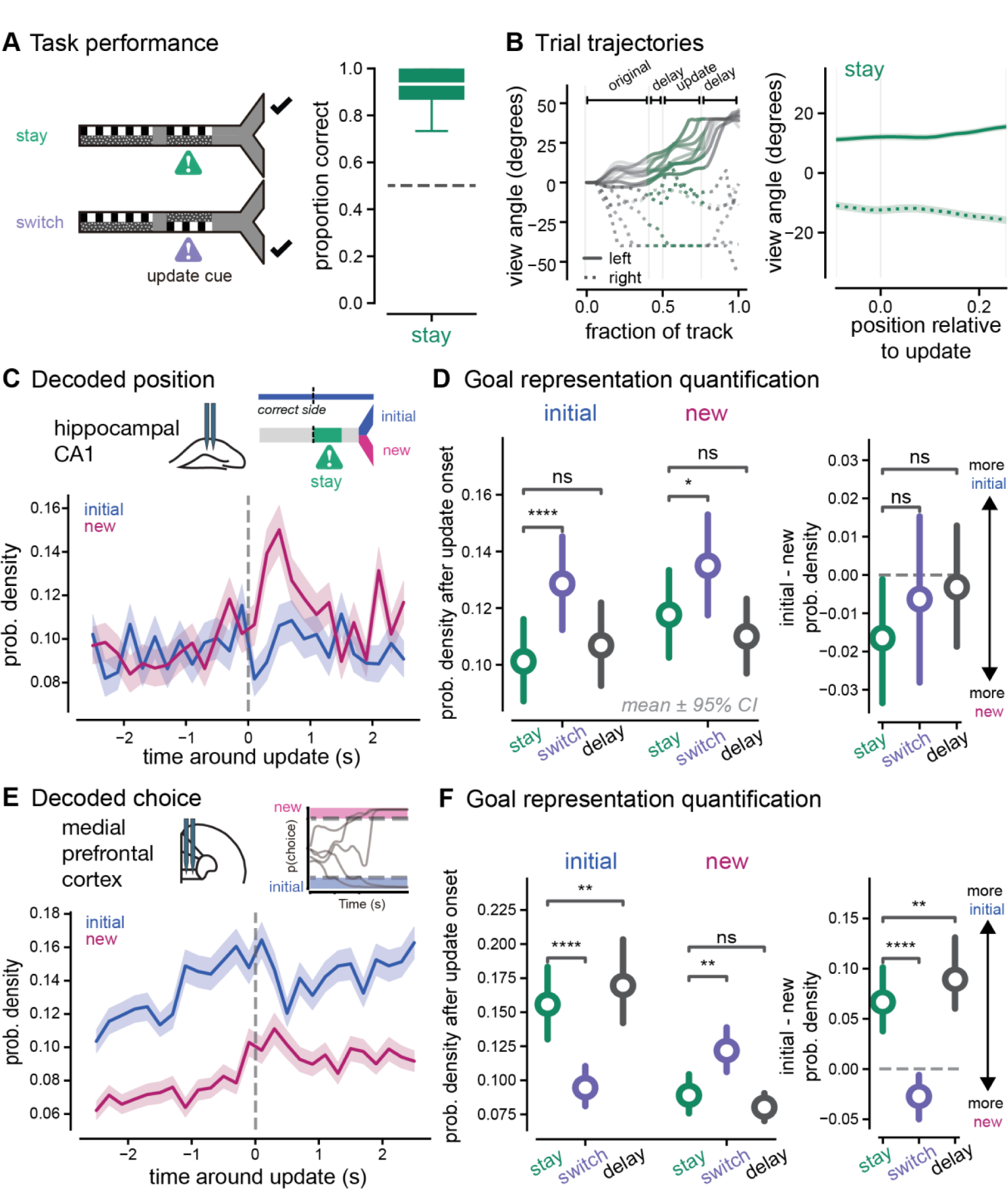
Prospective codes for updating behaviors are not due simply to new visual information **A.** *Left,* Schematic of stay trials in update task paradigm with cues displayed along the walls of the track. Stay trials (green, top, 10% of trials) consisted of an original cue period, a brief delay period, and then a second cue period during which the wall cues appeared on the *same* side as the original cues. Switch trials (purple, bottom, 25% of trials) were a similar structure, but during the second cue period the wall cues appeared on the *opposite* side as the original cues. Check marks indicate correct, rewarded side. *Right,* Proportion correct on stay trials (green) for all animals (n = 7 animals). Box plots indicate median and quartiles of distribution of 40-trial bins (stay: 0.92 ± 0.01, n = 117 windows, percentiles = 0.47, 0.87, 0.93, 1.00, 1.00). **B.** *Left*, Example behavioral trajectories for individual trials of each trial type. View angle (heading direction) across locations in the track for example correct left (solid lines) and right (dashed lines) trials. Task phases indicated with brackets and light grey vertical lines. Heading direction is set at 0 at trial onset. The delay period preceding the update location and the update cue location are shown in green. *Right,* Average view angle trajectories across all stay (green) trials. Dashed lines indicate initial right trials, solid lines indicate initial left trials, mean + SEM. **C.** *Top,* schematic of stay trial illustrating the update cue onset (green). On stay trials, a second visual cue is shown that indicates the initial arm is still correct. *Bottom,* probability densities of decoding the new (pink) and initial (blue) goal arms around the update cue on switch (center) and delay only (right) trials. Dark pink and blue lines indicate bins significantly different from baseline. Mean ± SEM across all trials shown. Note that the new arm is the correct arm on update trials but not on stay trials. **D.** Probability density decoding of the initial (left) and new (right) goal locations in the first 1.5 seconds after the update cue onset on stay (green), switch (purple), and delay only (black) trials. (initial, stay: 0.10 ± 0.00, n = 551 trials, percentiles = 0.00, 0.02, 0.07, 0.15, 0.58; new, stay: 0.12 ± 0.01, n = 551 trials, percentiles = 0.00, 0.03, 0.09, 0.18, 0.67;). Colored circle and line indicate mean ± 95% CI computed with n = 1000 bootstrap samples. *Right,* difference on single trials between initial and new probability densities after the update cue. (stay: -0.02 ± 0.01, n = 551 trials, percentiles = -0.66, -0.10, -0.01, 0.07, 0.51). Linear mixed effects model (LME) and ANOVA Tukey post-hoc, see Table 2 for statistical details. Descriptive statistics for switch and delay only trials can be found in the Figure 2 caption. **E.** *Top*, schematic of choice estimates. *Bottom*, integrated probability densities of decoding the new (pink) and initial (blue) choice estimates around the update cue on stay trials. Mean ± SEM across all trials shown. **F.** Probability density of decoding initial (left) and new (right) choices after the update cue on stay (green), switch (purple), and delay only (black) trials. (initial, stay: 0.14 ± 0.01, n = 483 trials, percentiles = 0.00, 0.05, 0.10, 0.20, 0.70; new, stay: 0.10 ± 0.00, n = 483 trials, percentiles = 0.00, 0.02, 0.07, 0.13, 0.66). *Right*, Difference between initial and new probability densities after the update cue (stay: 0.05 ± 0.01, n = 483 trials, percentiles = -0.66, -0.04, 0.03, 0.13, 0.70). Linear mixed effects model (LME) and ANOVA Tukey post-hoc, see Table 2 for statistical details. Descriptive statistics for switch and delay only trials can be found in the Figure 2 caption. ns, not significant, *p < 0.05, **p < 0.01, ***p < 0.001, ****p < 0.0001. All values reported as mean ± SEM, percentiles represent [minimum, 25^th^, median, 75^th^, maximum].

### Goal codes predict ability to accurately update decisions in response to new information

After finding that new, pivotal information increased non-local coding for both goal locations in hippocampus and caused a rapid switch from old to new choice codes in the prefrontal cortex, we then asked if these coding adaptations fail when animals do not accurately update decisions. We hypothesized that if increases in non-local goal codes in hippocampus contribute to flexible adaptation of navigational plans, then failure to switch goals in response to new information would co-occur with a failure to increase both non-local goal representations after the update cue is presented. To test this, we compared our population decoding of position in hippocampus and choice in prefrontal cortex on correct versus incorrect switch trials. To account for differences in speed (rotational and translational) as an explanatory variable for the amount of initial and new choice representations, we applied a general linear model (GLM). The residual goal codes from the model were then analyzed to assess changes in goal coding not explained by differences in speed and rotation (**Supplementary** Figure 9). We found that indeed on incorrect trials in hippocampus, the new goal choice coding was not as elevated as correct trials (p = 0.0025, new correct vs. incorrect, **Figure 6A**). However, the difference in new goal choice coding was small and highly variable. There was also no significant difference in initial goal choice coding on correct versus incorrect trials after the update cue (p = 0.2967 initial correct vs. incorrect, **Figure 6A**). Interestingly, *preceding* the update cue there was a noticeable increase in the initial goal arm representation that could not be explained by motor differences alone (p = 0.002, initial correct vs. incorrect). These results show that when animals failed to switch goals, in hippocampus new goal arm codes did not increase as much in response to the update cue, and initial goal arm codes were elevated *before* the update cue compared to correct trials.

**Figure 6.**
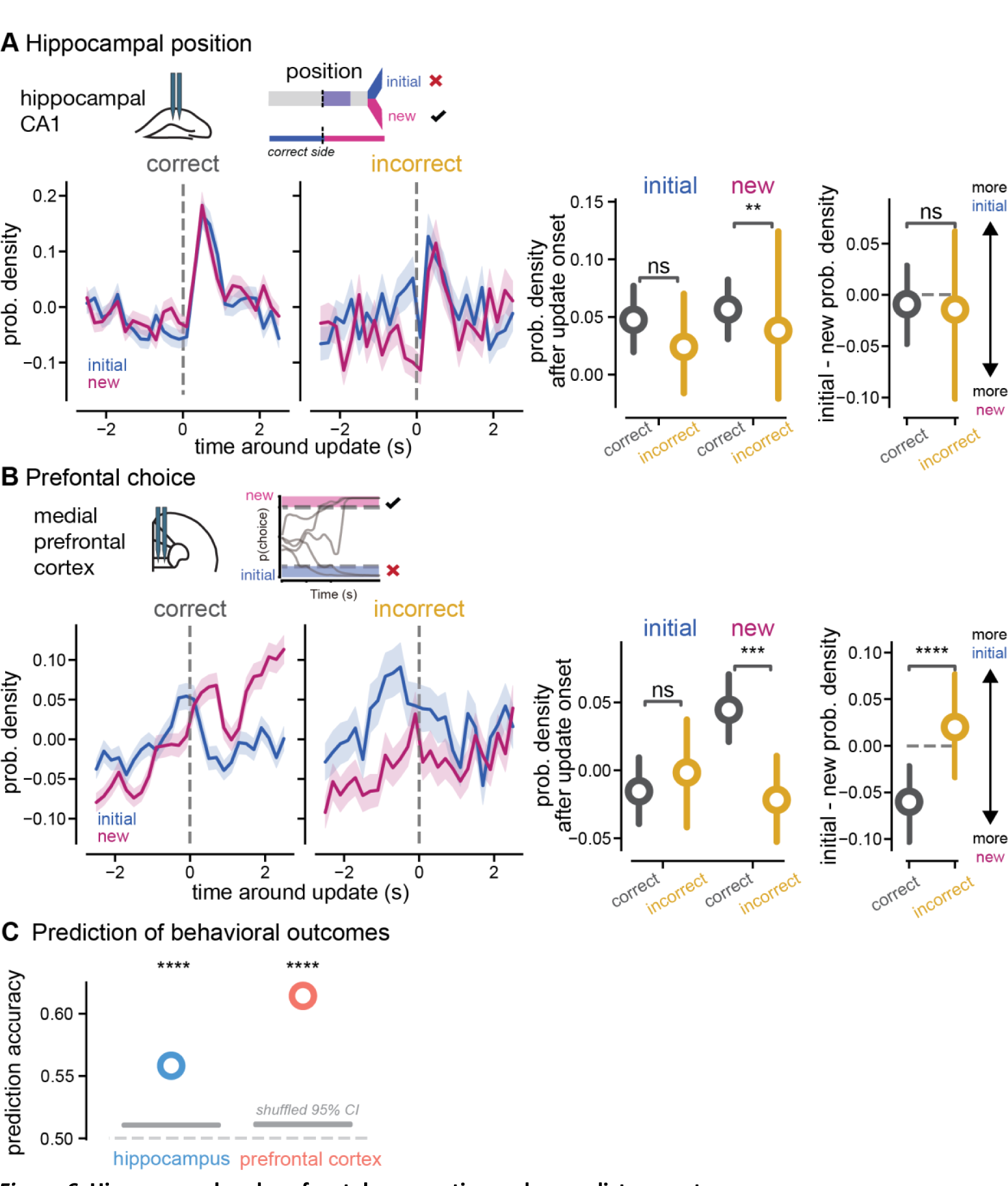
Hippocampal and prefrontal prospective codes predict correct responses. **A.** *Left*, residual decoding output (probability densities) from hippocampus of the new (pink) and initial (blue) position representations around the update cue on correct versus incorrect switch trials. Mean ± SEM across all trials shown. *Middle,* quantification of residual probability density differences after the update cue of hippocampal position codes on correct versus incorrect trials (black: correct, yellow: incorrect). Larger colored points and line indicate mean ± 95% CI computed with n = 1000 bootstrap samples (initial, correct: 0.06 ± 0.01, n = 1295 trials, percentiles = -0.50, - 0.21, -0.04, 0.22, 2.09; initial, incorrect: 0.03 ± 0.02, n = 407 trials, percentiles = -0.43, -0.22, -0.04, 0.18, 1.88; new, correct: 0.06 ± 0.01, n = 1295 trials, percentiles = -0.47, -0.20, -0.03, 0.25, 1.72; new, incorrect: -0.00 ± 0.02, n = 407 trials, percentiles = -0.45, -0.24, -0.08, 0.13, 2.27)*. Right*, quantification of difference between initial and new probability densities after the update cue (correct: -0.01 ± 0.01, n = 1295 trials, percentiles = -1.87, -0.29, -0.01, 0.27, 2.33; incorrect: 0.03 ± 0.02, n = 407 trials, percentiles = -2.49, -0.18, 0.03, 0.27, 1.87). Linear mixed effects model (LME) and ANOVA Tukey post-hoc, see Table 2 for statistical details. **B.** As in **A** for choice representations in prefrontal cortex. *Left*, residual integrated probability densities of the new (pink) and initial (blue) choice estimates around the update cue on switch (center) and delay only (right) trials. Mean ± SEM across all trials shown. *Middle:* quantification of residual probability density differences after the update cue of prefrontal choice codes on correct versus incorrect trials (initial, correct: -0.01 ± 0.01, n = 1109 trials, percentiles = -0.44, -0.19, -0.06, 0.10, 3.45; initial, incorrect: 0.01 ± 0.02, n = 359 trials, percentiles = -0.43, -0.18, -0.04, 0.12, 1.43; new, correct: 0.04 ± 0.01, n = 1109 trials, percentiles = -0.40, -0.15, -0.02, 0.17, 1.81; new, incorrect: -0.03 ± 0.01, n = 359 trials, percentiles = -0.42, -0.20, -0.10, 0.09, 1.61). *Right*, quantification of difference between initial and new probability densities after the update cue (correct: -0.05 ± 0.01, n = 1109 trials, percentiles = -2.03, -0.25, -0.04, 0.15, 3.62; incorrect: 0.04 ± 0.02, n = 359 trials, percentiles = -1.77, -0.18, 0.03, 0.22, 1.66). Linear mixed effects model (LME) and ANOVA Tukey post-hoc, see Table 2 for statistical details. **C.** Prediction of behavioral outcome (correct vs. incorrect final choice) using decoded choice estimates in prefrontal cortex and decoded position in hippocampus (55.83% accuracy for hippocampus and 61.43% accuracy for prefrontal cortex). Dashed lines indicate 50% prediction accuracy, grey lines indicated 95% CI for shuffled predictions. Permutation test, see Table 2 for statistical details. ns, not significant, *p < 0.05, **p < 0.01, ***p < 0.001, ****p < 0.0001. All values reported as mean ± SEM, percentiles represent [minimum, 25^th^, median, 75^th^, maximum].

We hypothesized that when animals failed to correctly respond to new information, the new and initial choice representations in prefrontal cortex would fail to flip from the initial choice being more strongly represented to the new choice being more represented. Indeed, we found new choice decoding did not overtake initial choice decoding on incorrect trials (**Figure 6B**). This lack of switching choice representations was driven by significantly smaller decoding of the new choice in prefrontal cortex on incorrect trials (p = 0.002, new correct vs. incorrect). In contrast, there was no significant difference in initial choice coding on correct and incorrect trials when accounting for motor differences (p = 0.5874, initial correct vs. incorrect, **Figure 6B**). Similar to hippocampus, the initial choice coding preceding the update cue was larger on average on incorrect versus correct trials, however in prefrontal cortex these pre-cue differences were not statistically significant when accounting for motor differences (p = 0.0536, initial correct vs. incorrect preceding update cue). Comparing new and initial choice representations within single trials, on incorrect trials the old and new choices were decoded at similar amounts even after new information was presented, in contrast to the significantly stronger new choice coding on correct trials (p < 0.0001, initial – new for correct vs. incorrect, p = 0.3531 initial – new vs. zero, incorrect trials, p = 0.0113 initial – new vs. zero, correct trials). These results show that when animals failed to switch destinations, initial choice representations decreased in response to new information similarly to correct trials, but there was a failure of the new choice codes to increase in prefrontal cortex after the update cue onset.

To test whether these observed codes were related to the animals’ behavioral performance, we then used the population measures of goal coding to predict behavioral outcomes on a trial-by-trial basis. We found that in both hippocampus and prefrontal cortex, the population representation of the initial and new goals after the update cue was presented predicted whether the animals would choose the correct or incorrect side (hippocampus: p < 0.0001, prefrontal cortex: p < 0.0001, permutation test, **Figure 6C**). It is important to note that the prediction accuracy was lower using hippocampal position decoding (55.83% accuracy) and slightly higher with prefrontal choice decoding (61.43% accuracy). Overall, we found hippocampal and prefrontal goal representations after the update cue onset predicted whether animals flexibly and accurately adapt plans on a trial-by-trial basis as would be expected if these codes were important to the animals’ behavioral performance. Together, these results show that when animals fail to adapt to new information, prefrontal represetantations of the new choice are lacking.

### Increased goal representations are correlated with choice commitment

Given that new information triggers a swift increase in hippocampal codes of possible goals and a rapid switch in prefrontal cortex from initial to new choice, we wondered if these changes are directly related to the amount animals must revise their ongoing behavior. We hypothesized that commitment to the initial decision at the time of the update cue onset would influence the degree of modulation of the initial and new goals representations. In other words, if the animals were less committed to their initial choice, the need to consider the alternative goal would be lower compared to trials where they were more committed, and more behavioral adaptation would be needed. To test this, we leveraged a behavioral readout of evolving choice in the virtual reality environment to separate trials with weak or strong commitment to the initial choice. This initial choice commitment was based on view angle at the time the update cue, with the top quartile of view angles (positive values pointing towards the initial goal arm) indicating strong commitment to the initial goal (**Figure 7A**). In hippocampus, when the animals were more committed to the *initial* side, the introduction of new, pivotal information resulted in an increased representation of the *new, alternative* side compared to trials where the animals were less committed or already heading towards the new side. Specifically, the view angle preceding new information was positively correlated with the increase in non-local goal representation of the new arm (p < 0.0001, r_s_ = 0.1540, new goal codes vs. view angle, **Figure 7B, Supplementary** Figure 10E). It is important to note that the correlation between view angle and prospective codes was only observed for the new arm and not the old arm, suggesting that this relationship is neural content-specific, not simply a general response due to visual cue differences at the time of the update (p = 0.3603, rs = 0.0261, initial goal codes vs. view angle, **Figure 7B**). Neural choice representations of the initial choice in prefrontal cortex covaried with view angle at the time of the update cue; if the animals were pointed more towards the initial side, the initial choice coding was higher than trials where the animals were pointed down the center or to the new side (p < 0.0001, r_s_ = 0.3344, initial choice codes vs. view angle, **Figure 7C**). Overall, these results demonstrate that greater commitment to the initial choice preceding new information leads to a larger increase of hippocampal non-local coding for the new side in hippocampus. On trials where animals need a larger trajectory adaptation in response to new information, non-local codes for the new goal destination are more prominent. Thus, while prospective codes in hippocampus represent both possible goals, the prominence of a particular goal representation depends on the degree of behavioral change needed.

**Figure 7.**
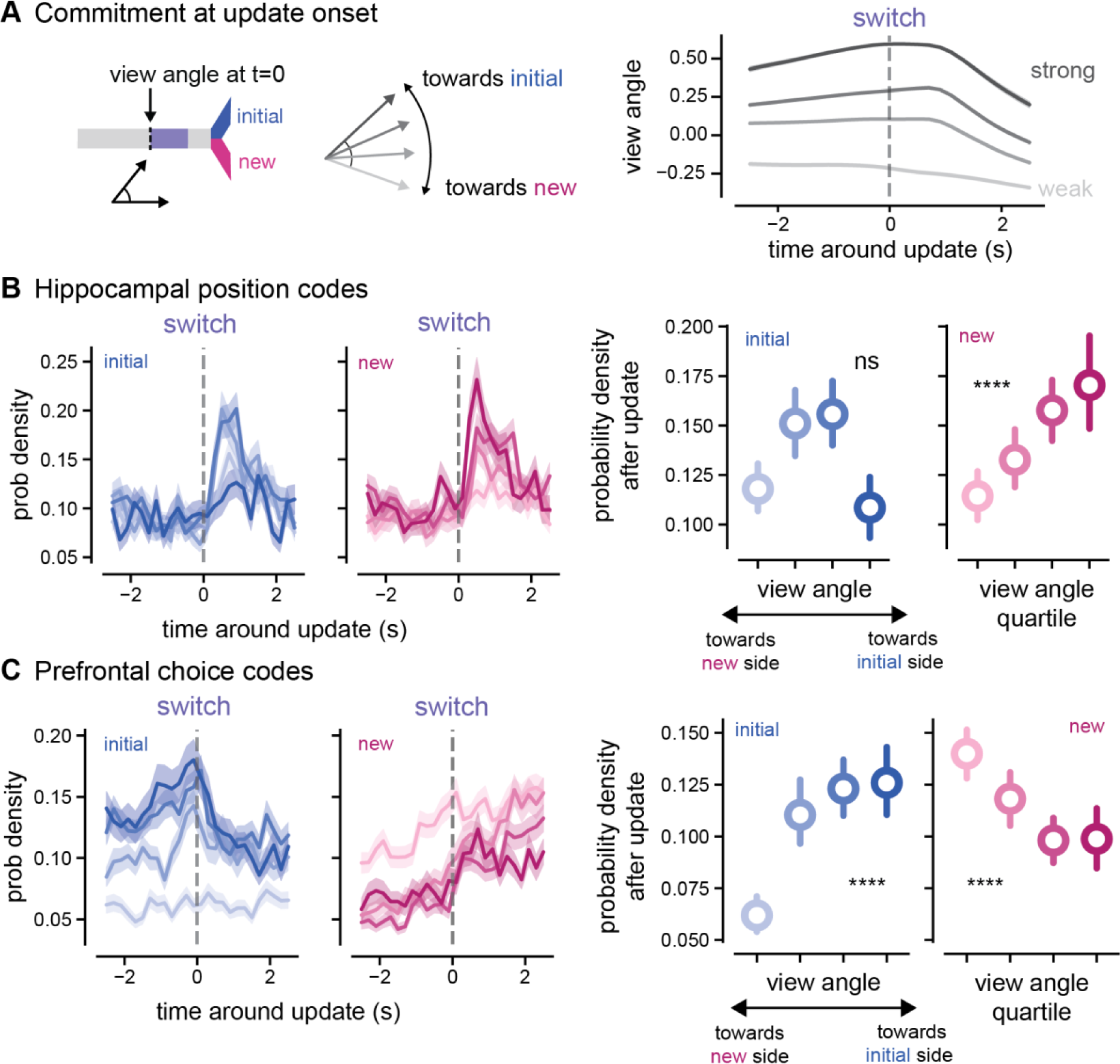
Hippocampal codes for *new* goal locations increase more strongly when animals are more committed to the initial goal. **A.** *Left*, schematic of commitment estimate breakdown from behavioral readouts around the update cue. *Right*, average view angle trajectories for all animals separated out into quartiles from greatest to least choice commitment at the time of the update cue onset. Darker indicates a stronger commitment to the initial side and lighter indicates a weaker commitment to the initial side. Mean ± SEM across all trials shown, note that SEM was very small. **B.** Hippocampal decoding of goal locations around the update cue onset on switch trials broken down by commitment quartiles based on the animals’ view angle at the time of the update cue onset on switch trials (shown in **A**). *Far left*, initial goal representations with darker blue indicating trials with a stronger commitment to the initial side measured by a greater view angle towards the initial side at update cue onset, and lighter blue indicating weaker commitment to the initial side, measured by a view angle towards the new side at update cue onset. *Center left*, as on far left for new goal representations (pink). *Right*, average probability density of decoding the initial goal (blue) or new goal (pink) locations in first 1.5 seconds after the update cue across all view angle quartiles (Initial goal representations by view angle quartiles, quartile 1 (towards new side): 0.12 ± 0.01, n = 410 trials, percentiles = 0.00, 0.02, 0.09, 0.18, 0.87; quartile 2: 0.15 ± 0.01, n = 305 trials, percentiles = 0.00, 0.03, 0.11, 0.23, 0.73; quartile 3: 0.16 ± 0.01, n = 324 trials, percentiles = 0.00, 0.05, 0.12, 0.23, 0.74; quartile 4 (towards initial side): 0.11 ± 0.01, n = 196 trials, percentiles = 0.00, 0.02, 0.09, 0.15, 0.56; New goal representations by view angle quartiles, quartile 1 (towards new side): 0.11 ± 0.01, n = 410 trials, percentiles = 0.00, 0.02, 0.07, 0.16, 0.83; quartile 2: 0.13 ± 0.01, n = 305 trials, percentiles = 0.00, 0.03, 0.09, 0.19, 0.90; quartile 3: 0.16 ± 0.01, n = 324 trials, percentiles = 0.00, 0.05, 0.12, 0.25, 0.94; quartile 4 (towards initial side): 0.17 ± 0.01, n = 196 trials, percentiles = 0.00, 0.05, 0.13, 0.24, 0.97). Spearman correlation, see Table 2 for statistical details. **C.** As in **B** for prefrontal neural codes for choice representation for initial (blue) and new (pink) choices (Initial choice representations by view angle quartiles, quartile 1 (towards new side): 0.06 ± 0.00, n = 357 trials, percentiles = 0.00, 0.01, 0.04, 0.09, 0.83; quartile 2: 0.11 ± 0.01, n = 270 trials, percentiles = 0.00, 0.03, 0.07, 0.14, 0.80; quartile 3: 0.12 ± 0.01, n = 286 trials, percentiles = 0.00, 0.05, 0.09, 0.16, 0.83; quartile 4 (towards initial side): 0.13 ± 0.01, n = 190 trials, percentiles = 0.00, 0.07, 0.10, 0.15, 0.85; New choice representations by view angle quartiles, quartile 1 (towards new side): 0.14 ± 0.01, n = 357 trials, percentiles = 0.00, 0.05, 0.11, 0.21, 0.62; quartile 2: 0.12 ± 0.01, n = 270 trials, percentiles = 0.00, 0.03, 0.09, 0.16, 0.50; quartile 3: 0.10 ± 0.01, n = 286 trials, percentiles = 0.00, 0.03, 0.07, 0.14, 0.69; quartile 4 (towards initial side): 0.10 ± 0.01, n = 190 trials, percentiles = 0.00, 0.03, 0.07, 0.13, 0.49). Spearman correlation, see Table 2 for statistical details. ns, not significant, *p < 0.05, **p < 0.01, ***p < 0.001, ****p < 0.0001. All values reported as mean ± SEM, percentiles represent [minimum, 25^th^, median, 75^th^, maximum].

## DISCUSSION

Using a navigation task with precise timing of environmental changes, simultaneous hippocampal-prefrontal cortex recordings, and decoding of position and choice, we determined how prospective codes change in response to new information to guide flexible decision-making. In our paradigm, animals rapidly update their behavior in dynamic environments in response to new information. In hippocampus, our results show that when new, crucial information is presented that requires animals to update their goal destinations, hippocampus increases non-local coding of *both* potential goal locations. Non-local goal representations in hippocampus increased more when animals needed to produce a larger trajectory change. In prefrontal cortex we found new, pivotal information causes choice codes to rapidly switch from representing old to new choices, prior to behavioral responses and prior to choice switching in hippocampus. On incorrect trials, hippocampal non-local goal codes remain relatively intact, but new prefrontal choice coding fails to increase. Together, these results show that non-local codes are generated in response to new, crucial information that requires flexible decision-making. New information triggers a rapid adaptation of ongoing plans, with hippocampus providing a swift increase in codes of possible goals and prefrontal cortex suppressing initial choices to select a new action plan. When more behavioral adaptation is needed, these prospective codes for new goals are stronger.

Questions about neural correlates of deliberation and planning are often posed within static environments in which behavioral planning is self-driven and internally modulated. However, one of the scenarios in which these codes might be used is rapid adaptation in response to a changing environment. Thus, our central question was how prospective codes in hippocampus and prefrontal cortex contribute to flexible decision-making in dynamic environments. Our results show that in hippocampus, non-local coding for goal locations increases when an environment requires a flexible change in action plans. Interestingly, in our task we observe that representations of *both* goal locations increase similarly in response to new information on correct trials. We did not observe that the planned goal was more strongly decoded than the alternative goal on correct trials until the animals reached the goal arms, tying in with previous work that action plans in rodents are not necessarily predicted by hippocampal prospective codes (Gillespie et al., 2021; Kay et al., 2020; Tang et al., 2021; M. Wang et al., 2020). We found some evidence of more subtle differences that might distinguish planned versus alternative choices in the hippocampus; e.g. significant theta modulation of new goal codes compared to non-significant modulation of initial goal codes. However, overall these results show that hippocampal prospective coding does not preferentially overrepresent the planned trajectory, but a general increase in prospective coding occurs when new, crucial information is presented in an environment.

Importantly, non-local goal coding of the new, upcoming goal increases more when animals need to produce a larger trajectory change in response to the new information. Our findings show that the representation of new possible goals in hippocampus reflects the animals’ need to consider new information in relationship to their current decision. These findings may explain variation in the ability to predict upcoming choices from hippocampal activity across studies. We propose that the non-local spiking events might reflect moments of representation of possible outcomes to the animals at a faster timescale scale than other noted behavioral deliberation events such as vicarious trial and error (Redish, 2016; Yu & Frank, 2021). While animals decrease their velocity slightly after the update cue is presented, the animals do not often pause at the update cue or choice point on the virtual reality track and thus do not exhibit traditional vicarious trial and error behavioral features during these moments of non-local coding. These non-local codes might be triggered not only by new information about goals, but also by task-relevant information more generally. Recent work found that neural codes in hippocampus of bats navigating down a tunnel shift to code for distance to other bats that might cause collisions or prompt changes in their trajectory (Sarel et al., 2022). This detection of relevant non-local stimuli results in a brief decrease in local position representations, similar to our findings. Our results show that non-local coding in hippocampus increases in response to task demands that require planning and flexibility, reflecting the need to consider new information and change behavior.

During flexible decision-making, we find that prefrontal cortex rapidly shifts from one choice to the other, suppressing the initial choice for the new choice to take over. Interestingly, prefrontal cortex choice switching occurs faster than that in hippocampus, suggesting that prefrontal cortex reaches a choice consensus more quickly. Prefrontal choice representational changes also preceded the animals’ turning movements on average, though these choice alterations may include some aspects of motor planning. Furthermore, when animals fail to flexibly change their goal destination in response to new information, prefrontal cortex choice representations look very different. In our task, we observe that when the new choice representation fails to overtake the old choice, animals are unable to flexibly respond to changes in the environment. This failure is likely occurring in prefrontal cortex; while non-local codes were disrupted in both brain regions on incorrect trials, hippocampal differences were more variable and had lower predictive power on trial outcomes than prefrontal cortex. However, this study is limited overall in that it does not address how necessary these neural signals from each brain region might be to the animals’ behavior. Instead, we manipulated the virtual reality environment to test how sensory information contributes to non-local coding and its role in decision-making. These results suggest that during decision-making in a dynamic environment, prefrontal cortex activity reflects a neural correlate of flexibility in the face of new information.

In summary, this work shows that prospective codes in hippocampus and prefrontal cortex are triggered in response to new, pivotal information in the environment to flexibly integrate new information to adapt navigational plans. We found that new, pivotal information causes rapid increases in prospective coding for non-local positions and switches in coding for upcoming choices and that these codes fail when animals are unable to adapt their previous choices. Furthermore, prospective codes for new goals are more strongly triggered when more navigation adaptation is needed. Thus, by examining neural responses to precisely-timed new information and the degree to which animals adapt to such information, our study provides unprecedented precision in addressing long-standing debates about how prospective codes support deliberation and planning. We propose these non-local codes are neural correlates of planning and consideration of alternative choices that can occur at faster timescales than observable behavioral deliberation and drive neural representations momentarily away from local coding when animals need to adapt their decisions and reconsider potential choices.

## METHODS

### Animals

All animal work was approved by the Institutional Animal Care and Use Committee at the Georgia Institute of Technology. Eight-week-old male *WT* mice on a C57Bl/6 background were obtained from the Jackson laboratory. Mice were single-housed on a reverse 12-hour light/12-hour dark cycle. At the start of the behavioral and electrophysiological experiments, mice were food-restricted to 85% percent of their baseline body weight, and water was provided without restriction. Fifteen animals total were trained for electrophysiological recordings, one mouse was excluded for lack of movement and inability to complete trials in the visually guided phase, seven were excluded for never reaching the final phase of the task (usually due to difficulties in performing over longer delays), and one was excluded after electrophysiological recordings were started due to lack of performance during recording sessions, resulting in seven mice total in the final dataset.

### Surgical procedures

Animals were implanted with headplates at approximately 8 weeks of age. Mice were anesthetized with isoflurane before headplate implant surgery. A custom stainless steel headplate was fixed to the skull using dental cement (C&B Metabond, Parkell), and the target craniotomy site for recordings was marked on the skull (in mm, from bregma: −2.0 anterior/ posterior, +/-1.8 medial/lateral for hippocampal CA1 and +1.0 anterior/posterior, +/- 1.25 medial/lateral for medial prefrontal cortex). Craniotomies were later performed before electrophysiology recording sessions in mice that reached the final phase of the task. These craniotomies (200-500um diameter) were made using a dental drill to thin the skull and then opening a small hole in the skull with a 27-gauge needle. Craniotomies were sealed with a sterile silicone elastomer (Kwik-Sil WPI) and only opened for recording experiments.

### Behavioral task and training

The virtual reality environment was designed using ViRMEn (Aronov & Tank, 2014) software in Matlab and displayed on a cylindrical screen using a projector system reflected by two mirrors. Head-fixed animals ran on a spherical treadmill composed of an 8-inch polystyrene foam ball floating on air. The ball movement was recorded with optical mice and converted to velocity signals in LabView. Pitch velocity was used for forward motion through the environment, and roll velocity was used for rotational velocity. The view angle was defined as the direction the animals were facing in the virtual reality environment. The start of the trial had a view angle of 0 degrees when the animals were facing straight ahead. View angles began to veer positive (towards the right) or negative (towards the left) as the rotational and translational velocity inputs from the animals’ movement on the ball were translated into movement in the virtual reality space. Rewards of sweetened condensed milk (1:2 dilution in water) were delivered via a reward spout and licks were detected using a photo-interrupter.

The behavioral task was a virtual y-maze in which animals had to choose to go to the left or right to receive a reward. The full length of the maze was approximately 3 m for the center arm. Trials were structured as follows: first, the screen displayed the central arm of the environment, and the mouse’s position was frozen in place for 3 seconds to allow the mouse time to adjust its running patterns before it began to move down the track. After this 3s period, the mouse could begin moving forward, but the view angle and x-position were restricted to 40 degrees and 0 virtual displacement for the first 0.05 fraction of the track. Then, once the mouse passed the initial zone of the track, the mouse could rotate and move freely. The visual cues turned on and off at different locations in the track as the animals passed. On correct trials, a reward was delivered at the end of the track, the VR screen froze for 3 seconds, and then the task shifted into an intertrial interval period of 6 seconds. On incorrect trials, the task immediately shifted into a 12- second intertrial interval, which was longer as a form of punishment.

The animals underwent several phases of behavioral shaping and training to reach the final version of the task. In the first phase, animals ran down a linear track that was used to acclimate the mouse to the head-fixed VR setup. The linear track increased in length until the animals completed a certain number of trials for each track length. The second phase of the task was a short y-maze choice task where visual cues indicating the correct and incorrect locations were visible for the full length of the track. After the animals performed two sessions above 75 percent correct, they advanced to the long y-maze choice task with the same visually guided trials as the previous phase but now with the central arm the same length as the final task. Animals were then advanced through phases with a delay at the end of the track. During the delay, the visual cues indicating the correct left or right side were no longer visible and the walls of the track were grey instead. This delay was moved gradually earlier in the track, making it longer in time, through three separate phases. After the animals had reached the learning criteria in the third delay phase, the update cue trials were introduced. In these trials, a second visual cue appeared after the first original visual cue, indicating either the same side as the initial cue or the opposite side. Once the animals showed signs of understanding these second cues, the delay location was moved earlier once more, so that the final update task trial structure had three different trial types. These trial types consisted of delay only trials (65% of trials), in which the initial cue was shown and there was a long delay until the end of the track, switch trials, in which there was an original cue, brief delay, and second cue on the opposite side of the track as the original cue (25% of trials) and finally stay trials, in which there was an original cue, brief delay, and second cue on the same side of the track as the original cue (10% of trials). To control for side biases, the animals’ preference for the left or ride side of the track, we applied a bias correction algorithm for selecting the next trial type. In short, the probability that the next trial would be on the left or right side was dependent on the history of left and right trial errors. If the animals preferred turning to the right arm at the choice point and had missed mainly left trials, then there would be a higher likelihood of the next trial being on the left, or non-preferred side (Hu et al., 2009; Pinto et al., 2018). This algorithm was applied throughout training and during electrophysiological recordings. Animals were trained approximately 5-7 days per week, 1 hour per day on average. Overall, this training process took 55.43 ± 7.38 days on average (mean ± SEM, n = 7 animals).

### Electrophysiological recordings

Recordings occurred during behavioral task performance when animals navigated through the virtual reality environment. Data were acquired using a SpikeGadgets acquisition system with a sampling rate of 30kHz and a ground pellet as reference. Animals were head-fixed on the treadmill for a maximum of one five-hour-long recording session per day (number of sessions ranged from 6-12 per animal, see **Table 1** for details). A 64-channel, dual-shank NeuroNexus probe was placed in a slightly different location within the craniotomy at the beginning of each recording session and advanced to the target location with the angles of the manipulators adjusted according to the final craniotomy location and target location. In hippocampus, the target location was -1.8-2.0 anterior/posterior, 1.5-1.8 medial/lateral, and 1.4 dorsal/ventral. In medial prefrontal cortex, the target location was 1.7-1.8 anterior/posterior, 0.4 medial/lateral, and 2.0 dorsal/ventral. For hippocampal recordings, the probe was advanced to the CA1 pyramidal layer of hippocampus identified via electrophysiological characteristics: large theta waves, sharp-wave ripples, and 150+ μV spikes on multiple channels. Recording sites usually spanned the layer. For medial prefrontal cortex recordings, the probe was advanced to the target location using the distance traveled as the primary metric, until a suitable location was found with the maximum number of spikes.

During the final recording session from each hemisphere, a probe was coated with DiI and inserted to the target depth. Brains were then drop-fixed in 4% paraformaldehyde. Brains were sectioned to either 100um thick with a Leica VT1000S vibratome or they were sectioned to 60um thick with a Leica cryostat after freezing at -80 degrees. Sections were then stained with 0.2% 1mMol DAPI and mounted on microscopy slides with Vectashield mounting medium. Images were acquired on a Zeiss confocal microscope. All images with visible DiI were registered to the Allen brain atlas using SHARP-Track (Shamash et al., 2018). Probe regions of interest were added based on DiI location, isolating individual shanks of the electrode when possible, and the software determined a best-fit line as an approximation of the probe path. The deepest identified point of visible DiI was used to calculate an estimated recording location in the Common Coordinate Framework.

### Local field potential and single unit preprocessing

The local field potential was obtained by downsampling raw traces to 2kHz and bandpass filtering between 1-300Hz. Outliers were eliminated by interpolating over outliers when the pre-filtered LFP signal was 15 standard deviations above the mean. All LFP analyses used the signal from a single channel that was putatively located in the *stratum pyramidale*. To identify this channel, the LFP was bandpass filtered for the sharp-wave ripple band (150-250 Hz, see details below) and the average of the sharp-wave ripple band envelope over time was calculated from each channel. The channel with the highest average sharp-wave ripple band power was used for all further LFP analyses, and this channel was predominately located in the middle of the depth-wise span of the NeuroNexus probe.

Spike detection and sorting were performed using Kilosort 2.0 spike sorting algorithm (Pachitariu et al., 2016) and then were manually curated using Phy software. Cell types were classified into putative pyramidal cells and narrow interneurons and wide interneurons using the default spike width and the autocorrelogram criteria from CellExplorer software (Petersen et al., 2021): narrow interneuron if the trough to peak value was <= 0.425 ms, wide interneuron if the trough to peak value was > 0.425 ms and the autocorrelogram tau rise value greater than 6 ms, and the remaining cells were classified as pyramidal cells. The classified distributions of neurons were compared to the ground-truth mouse data provided in the software as a visual confirmation of the accuracy of the classification. Positive spikes were identified with a polarity > 0.5 and were flipped for spike width calculations. All cells were visually inspected and a small number (less than 10 total) were excluded if the spike width could not be accurately quantified with the waveform.

### Behavioral analysis

To quantify behavioral performance, we binned behavioral data into 40 trial windows for each trial type to calculate a rolling window of proportion correct throughout the behavioral sessions. We also obtained the spatial trajectories of the animals by calculating an average value for each position bin in the environment across individual trials, and then averaging the left and right trials separately to obtain a final average trajectory for each behavioral readout.

For position representation analyses, the 2D position of the VR environment was converted into a 1D linearized position. Linearized position was calculated by generating a graph of the VR environment with 3 segments and 4 nodes: 1 node was at the start of the track, 1 node was at the choice point at the end of the central arm and between the two choice arms, and then the 2 final nodes were at the end of each choice arm. We used the track-linearization package to project each position in the environment using an HMM (hidden Markov map). Artificial gaps between the segments of the track were added for visualization purposes.

Locomotion periods were defined as times when the animals’ movement was above a velocity threshold. To calculate this threshold, we plotted the distribution of optical mouse recorded velocities and observed a bimodal distribution. We selected a threshold that separated these two distributions and used that for subsequent decoding analyses. Multiple thresholds were tested and the results remained similar.

### Choice modeling

To calculate an estimate of choice as animals ran through the environment, we built an LSTM neural network using TensorFlow based on previous choice estimate values obtained in virtual reality two-choice maze tasks (Tseng et al., 2022). The LSTM network contained a 10-unit LSTM layer, followed by a 1-unit dense layer with sigmoid activation to predict the reported choice at every time point in the trial using the velocities, view angle, and position from that trial up until that time point. The network was compiled with an Adam optimizer and a binary cross-entropy loss function. The input data was a matrix of the rotational and translational velocity signals, the forward position in the maze, and the view angle of the mouse at each time point. A separate model was trained for each behavioral session using a combination of model-averaging and k-fold cross-validation to train and test the model. We divided the data using stratified k-fold cross-validation with 6 folds, in which a separate model was trained for each group of 5 folds and used to predict/test on the 5^th^ fold. The final prediction for each trial was the average of three repeats of this 6-fold cross validation. The training data was normalized and padded to the length of the longest trial, but all trials with a length greater than twice the average were excluded. The hyperparameters of the neural network were selected using a grid search on a subset of the recording sessions and were as follows: batch size = 32, epochs = 20, learning rate = 0.1. Finally, we took the prediction of the model and calculated the log likelihood with log base 2 to assess the overall performance. The prediction data was then used as a feature input to the Bayesian decoding model to assess how the neural activity represented evolving choice as the animals run through the environment.

### Decoding analyses

To identify a neural representation of different features of the task at a population level, we performed a Bayesian decoding analysis to estimate the probability of different positions or choices given the observed spiking activity at that period. For all decoding analyses, we analyzed hippocampal CA1 and medial prefrontal cortex separately. We first excluded any sessions in which there were less than 20 single units for the brain region of interest and there were less than 50 trials for the behavioral session. These thresholds were chosen by visually inspecting and quantifying the overall decoding accuracy of each session and determining a minimum criteria in which high-fidelity decoding could be obtained. These criteria were applied separately to each brain region (e.g., data from one brain region might be included for the session if it met the criteria, but the other brain region would be excluded if it did not). Brain region decoding outputs were also calculated separately. To build our encoding model, we used only periods within a behavioral trial (excluding the intertrial interval) and only locomotion periods when the animal was moving. We used both correct and incorrect trials but only trials in which there was no update cue. Feature tuning curves were computed by calculating the number of spikes per feature bin (n = 50 bins for all features) and normalizing for occupancy time. For decoding, we used 25ms when analyzing decoding by individual theta phase and 200ms windows otherwise. To perform the decoding calculations, we used the Pynapple (Viejo et al., 2022) analysis package using a uniform prior. In a subset of our analyses, we used all trial types (delay only, switch, and stay trials) to build our encoding model as a control. To validate our decoding output, we built the encoding model using 80% of the data and held out a test set of 20% of the data to confirm decoding accuracy across the virtual reality environment.

To calculate representational changes around the update cue, we integrated the posterior probability densities for the initial choice arm and new choice arm based on the boundaries defined by our track graph used for track linearization. We calculated decoding error as the difference between the predicted and actual position. For some analyses, we converted the decoding output into terms of probability density / chance (Saleem et al., 2018; Sarel et al., 2022), where chance was defined as a uniform representation across all spatial bins. This probability density / chance value was obtained by multiplying the probability density function by a uniform constant of 1 / the number of spatial bins. With this output, a value of 1 indicates that the decoding output of that bin is the amount expected if all locations were uniformly represented and no location was represented more than others. A value equal to the number of spatial bins would indicate the maximum likelihood of the bin being represented.

To quantify the changes in prospective goal coding around the update cue, we calculated the average integrated decoding output value in the 0 to 1.5 second window after the update cue was presented to compare relative amounts of prospective coding between trial types. We also quantified the difference in initial and new goal representation on a trial-by-trial basis to compare relative amounts of initial and new goal representations between trial types. For analyses in which we quantified differences in goal coding preceding the update cue onset, we used the -1.5 to 0 second window as our baseline. When quantifying choice codes, we applied a 10% threshold to quantify strong choice representation of either the new or initial choice. We assessed other thresholds and found similar results.

To assess how choice commitment affected neural representations around the update cue, we obtained the view angle at the time of the update cue onset for each trial. We then split this view angle data into quartiles and with these separately grouped trial types, used the same time windows and quantification analysis described above to calculate the average responses.

### Theta cycles and phase

For each session, we identified a channel as the putative pyramidal channel in CA1 using sharp-wave ripple power as described above. This channel was used as our LFP recording site. To isolate hippocampal theta oscillations, the LFP was bandpass filtered for theta (4-12 Hz) using an FIR (finite impulse response) equiripple filter. Peaks and troughs of the filtered LFP were detected and used to define the phase of the LFP. In order to use a common theta reference across recording sessions, we calculated a phase histogram of all putative pyramidal cell spiking activity in CA1 for a recording session using 30-degree bins. We then adjusted the theta phase measurement to be 0 degrees at the location of maximal CA1 firing.

To calculate theta phase modulation of goal decoding, we calculated theta phase histograms by identifying the theta phase at the center time of each decoding window and then calculating the integrated posterior probability densities for each theta phase bin for each trial. To compare values across theta, we divided the theta oscillation into quarters and calculated the average decoding output value for each quarter of the theta oscillation.

### Generalized linear model (GLM) analysis

To account for the effects of speed and direction on the differences in neural codes between correct and incorrect trials, we applied a GLM with rotational and translational velocities as the explanatory variables or predictors and the goal decoding output as the response variable. A separate model was trained for the different goal representations, initial and new. To fit the model, we used the velocity and decoding output values in 200ms bins in the 2.5 seconds before and after the update cue onset across all trial types and animals. We then plotted the Pearson residuals as a measure of the decoding output not explained by speed and direction differences between trial types. Cox and Snell’s pseudo r-squared values were reported as an estimate of model fit.

### Prediction of behavioral choice

For prediction of animals’ final choice using neural activity, we performed a trial-by-trial classification analysis using support vector machines (SVMs). For each region and update trial type, we trained independent SVMs. For each trial, we used the quantification of goal representation difference between the initial and new goal as described above as a feature (n=1, initial – new) to predict the animals’ final choice (k = 2, correct or incorrect). Before training the classifier, we balanced the correct and incorrect trial classes using random resampling with replacement until both target classes had the same number of trials. We used a radial basis function (Gaussian) kernel for all SVMs, and we selected the hyperparameters (*C* and *γ)* using a random search method with leave one out cross-validation to prevent overfitting. After performing the random search over n=10 iterations to optimize classification accuracy, the best hyperparameters were selected. These hyperparameters were then used and a leave-one-out cross validation procedure was used to assess the overall accuracy of the classifier (percentage of trials correctly classified). This randomized hyperparameter search and final accuracy assessment were performed separately for each update trial type. As a control, the randomized search and cross validation were also performed with randomly shuffled target (choice) labels n=1000 times. The significance of the classifier was assessed by testing whether the classifier outperformed 95% of the distribution of accuracies from the shuffled classifier (permutation test).

### Statistical analyses

We used a linear mixed-effects models (LME) approach to assess the significance of differences while controlling for repeated measures from the same animals and sessions. In our model, our variables of interest such as decoding output or single unit firing rates were included as our fixed effects. We set our random effects as session-nested-in-animal random effects with random intercepts at both levels. Statistical significance was first estimated with an ANOVA with Kenward-Roger’s methods to determine whether the predictors had any significant effect. If the F-test was statistically significant, we then performed pairwise comparisons to assess significant differences using estimated marginal means and reported Tukey-adjusted p-values. For one-sample analyses, we used a Wilcoxon signed rank test. For correlation analyses, we calculated the Spearman rank-order correlation coefficient. All statistical tests were two-sided unless stated otherwise. Details on specific statistical parameters and the values of sample size n are described in the figure legends and statistical table (**Table 2**).

Analyses were performed using custom pipelines in Matlab, Python, and R with the following libraries: NumPy, SciPy, Matplotlib, Scikit-learn, Pandas, Tensorflow, Pynapple, nwbwidgets, pynwb, seaborn, track- linearization, pingouin, statannotations, statsmodels, lmer, lmerTest, emmeans, SHARP-Track, Kilosort, Phy.

## ACKNOWLEDGEMENTS

We thank Shantanu Jadhav for helpful feedback and all members of the Singer laboratory for technical assistance and feedback on the paper. We thank Bailey Mariner and scidraw.io for illustrations. This work was supported by NSF GRFP grant DGE-1444932, NIH grant T32NS096050-22, and the Emory University Women’s Club Graduate Research Fellowship (to SMP); the Georgia Institute of Technology President’s Undergraduate Research Fellowship (to NK and TAY); the National Institutes of Health (NIH)-National Institute of Neurological Disorders and Stroke Grant R01 NS109226, the NIH National Institute of Aging Grant RF1AG078736-01, Packard Award in Science and Engineering, McCamish Foundation, Friends and Alumni of Georgia Tech (to ACS).

## Author Contributions

SMP and ACS conceptualized the study and developed methodology. SMP, TAY, NK, and TCR carried out behavioral training. SMP conducted electrophysiological recordings. SMP and TAY curated data. SMP analyzed the data. SMP and ACS wrote the paper. ACS supervised the project and acquired funding.

## Data and Materials Availability

Custom code is available on GitHub: https://github.com/stephprince/update-project. All data used in this study will be shared on public repositories at the time of publication.

