## Supplementary Information for "New information triggers prospective codes to adapt for flexible navigation"

*Includes*

2 Tables

10 Figures

### SUPPLEMENTARY TABLES

| Animal | Total sessions | Medial prefrontal cortex approximate* locations | Hippocampal CA1 approximate* locations | Single units** | Behavioral trials*** |
|---|---|---|---|---|---|
| S17 | 8 | Prelimbic area layer 6a:<br>A/P: 2.02, D/V: 2.94, M/L: -0.75<br>A/P: 2.0, D/V: 2.89, M/L: 0.8,<br>Anterior cingulate area dorsal part layer 6a:<br>A/P: 2.02, D/V: 2.52, M/L: -0.77<br>A/P: 1.91, D/V :2.64, M/L: 0.99 | A/P: -1.8, D/V: 1.7, M/L: 1.68<br>A/P: -2.1, D/V: 1.87, M/L: -1.37<br>A/P: -2.1, D/V: 1.84, M/L: -1.49 | Total: 646<br>mPFC: 247<br>CA1: 399 | Total: 591<br>Switch: 114<br>Stay: 48<br>Delay: 429 |
| S20 | 12 | Anterior cingulate area ventral part layer 5:<br>A/P: 1.38, D/V: 3.41, M/L: -0.38<br>A/P: 1.36, D/V: 2.68, M/L: -0.58 | A/P: -2.45, D/V: 1.63, M/L: -1.26<br>A/P: -2.46, D/V: 1.6, M/L: -1.35 | Total: 977<br>mPFC: 272<br>CA1: 705 | Total: 1689<br>Switch: 303<br>Stay: 117<br>Delay: 1269 |
| S25 | 10 | Prelimbic area layer 6a:<br>A/P: 1.63, D/V: 3.05, M/L: -0.74<br>Prelimbic area layer 2/3:<br>A/P: 1.62, D/V: 2.89, M/L: 0.25<br>Prelimbic area layer 1:<br>A/P: 1.62, D/V: 2.74, M/L: 0.15 | A/P: -2.12, D/V: 1.53, M/L: 1.49<br>A/P: -2.13, D/V: 1.4, M/L: 1.59 | Total: 906<br>mPFC: 286<br>CA1: 620 | Total: 2244<br>Switch: 421<br>Stay: 189<br>Delay: 1634 |
| S28 | 11 | Slices damaged | Slices damaged | Total: 1155<br>mPFC: 550<br>CA1: 605 | Total: 2137<br>Switch: 344<br>Stay: 108<br>Delay: 1685 |
| S29 | 12 | Slices damaged | A/P: -1.78, D/V: 1.78, M/L: -1.5<br>A/P: -1.78, D/V: 1.72, M/L: -1.58<br>A/P: -1.52, D/V: 1.72, M/L: 1.37<br>A/P: -1.55, D/V: 1.57, M/L: 1.59 | Total: 1530<br>mPFC: 715<br>CA1: 815 | Total: 2308<br>Switch: 422<br>Stay: 161<br>Delay: 1725 |
| S33 | 6 | Infralimbic area layer 6a:<br>A/P: 1.74, D/V: 3.49, M/L: -0.59<br>Infralimbic area layer 5:<br>A/P: 1.74, D/V: 3.45, M/L: -0.51 | A/P: -2.32, D/V: 1.51, M/L: -1.66 | Total: 617<br>mPFC: 178<br>CA1: 439 | Total: 696<br>Switch: 97<br>Stay: 42<br>Delay: 557 |
| S34 | 7 | Infralimbic area layer 6a:<br>A/P: 1.81, D/V: 3.79, M/L: 0.77<br>A/P: 1.77, D/V: 3.66, M/L: 0.6l<br>A/P: 1.8, D/V: 3.73, M/L: -0.57<br>A/P: 1.83, D/V: 3.46, M/L: -0.69 | A/P: -1.58, D/V: 1.61, M/L: -1.56<br>A/P: -1.65, D/V: 1.72, M/L: -1.48<br>A/P: -1.88, D/V: 1.58, M/L: 1.32<br>A/P: -1.88, D/V: 1.57, M/L: 1.55 | Total: 618<br>mPFC: 309<br>CA1: 309 | Total: 996<br>Switch: 139<br>Stay: 52<br>Delay: 805 |

***Table 1*** **Experimental metadata for all recording sessions.**

*Recording locations are approximate and indicate the DiI-identified locations registered to the Allen Common Coordinate Framework and mouse atlas from the last recording session of each animal for that hemisphere. Exact locations varied slightly between sessions due to day-to-day differences in electrode insertions. Hippocampal CA1 locations were also confirmed with electrophysiological signatures (see **Methods**).
**Number of units indicates final unit counts after spike sorting, quality curation, and cell type classification (see **Methods**).
***Number of trials indicate all completed trials during recording sessions.

| Panel | Data | Group size | Statistical method | Comparison | Test statistic | P-value | Notation |
|---|---|---|---|---|---|---|---|
| Figure *1*B | Proportion correct | N = 66 sessions N = 7 animals | Wilcoxon signed-rank test | Switch trials vs. 50% correct | W = 48.5 | <0.0001 | **** |
| | | | | Delay only trials vs. 50% correct | W = 125.5 | <0.0001 | **** |
| Figure *2*D | Hippocampal position decoding; posterior probability density integrated by goal arm | Switch: n = 1295 trials Delay only: n = 926 trials | Linear mixed effects model; ANOVA, Tukey post-hoc pairwise test | Trial type (switch, delay only), goal arm | F(4400.90) = 21.79 | < 0.0001 | **** |
| | | | | Switch vs. delay only, initial goal | Z = -5.87 | < 0.0001 | **** |
| | | | | Switch vs. delay only, new goal | Z = -5.39 | <0.0001 | **** |
| Figure *2*D | Hippocampal position decoding; difference in initial vs. new goal arm posterior probability density | Switch: n = 1295 trials Delay only: n = 926 trials | Linear mixed effects model; ANOVA | Trial type (switch, delay only) | F(2205.71) = 0.08 | 0.7806 | ns |
| Figure *2*D | Hippocampal position decoding; difference in initial vs. new | Switch: n = 56 sessions, Delay only: n = 56 sessions | Wilcoxon signed-rank test | Switch vs. zero | W = 676 | 0.3216 | ns |
| | | | | Delay only vs zero | W = 706 | 0.4554 | ns |
| Figure 2E | Prefrontal position decoding; posterior probability density integrated by goal arm | Switch: n = 1109 trials Delay only: n = 798 trials | Linear mixed effects model; ANOVA, Tukey post-hoc pairwise test | Trial type (switch, delay only), goal arm | F(3773.42) = 1.91 | 0.1261 | ns |
| | | | | Switch vs. delay only, initial goal | Z = -1.65 | 0.3496 | ns |
| | | | | Switch vs. delay only, new goal | Z = -1.72 | 0.3148 | ns |
| Figure 2E | Prefrontal position decoding; difference in initial vs. new goal arm posterior probability density | Switch: n = 1109 trials Delay only: n = 798 trials | Linear mixed effects model; ANOVA | Trial type (switch, delay only) | F(1903.52) = 0.012 | 0.9111 | ns |
| Figure 2E | Prefrontal position | Switch: n = 47 sessions, | | Switch vs. zero | W = 471 | 0.3227 | ns |

| | | | | | | | |
|---|---|---|---|---|---|---|---|
| | decoding; difference in initial vs. new | Delay only: n = 47 sessions | Wilcoxon signed-rank test | Delay only vs zero | W = 545 | 0.8448 | ns |
| Figure *3*B | Hippocampal position decoding; pre vs. post update cue onset; first half theta cycle | Switch: n = 1295 trials | Linear mixed effects model; ANOVA Tukey post-hoc pairwise test | Pre and post update, location | F(7709.65) = 142.33 | < 0.0001 | **** |
| | | | | Pre, central vs. post, central | Z = -16.06 | < 0.0001 | **** |
| | | | | Pre, initial vs. post, initial | Z = 10.28 | < 0.0001 | **** |
| | | | | Pre, new vs. post, new | Z = 9.91 | < 0.0001 | **** |
| Figure *3*C | Hippocampal position decoding; phase modulation by location | Switch: n = 1295 trials | Linear mixed effects model; ANOVA Tukey post-hoc pairwise test | Phase and location | F(7709.45) = 18.93 | < 0.0001 | **** |
| | | | | First half, initial vs. central | Z = -3.05 | 0.0275 | * |
| | | | | First half, new vs. central | Z = -3.35 | 0.0106 | * |
| | | | | Second half, initial vs. central | Z = 6.19 | < 0.0001 | **** |
| | | | | Second half, new vs. central | Z = 8.31 | < 0.0001 | **** |
| Figure *3*C | Hippocampal position decoding; phase modulation | Switch: n = 1295 trials | Linear mixed effects model | Central arm, first half vs. second half | F(2533.33) = 119.77 | < 0.0001 | **** |
| Figure *3*C | Hippocampal position decoding; phase modulation | Switch: n = 1295 trials | Linear mixed effects model | Initial arm, first half vs. second half | F(2533.36) = 0.65 | 0.4203 | ns |
| Figure *3*C | Hippocampal position decoding; phase modulation | Switch: n = 1295 trials | Linear mixed effects model | New arm, first half vs. second half | F(2533.71) = 11.22 | 0.0008 | *** |
| Figure *3*C | Hippocampal position decoding; theta timescale update trial types | Switch: n = 1295 trials; Stay: n = 552 trials; delay only: n = 927 trials | Linear mixed effects model; ANOVA Tukey post-hoc pairwise test | Trial type (switch, delay only), initial goal arm | F(5496.53) = 27.48 | < 0.0001 | **** |
| | | | | Switch vs. delay only, first half | Z = -5.79 | < 0.0001 | **** |
| | | | | Switch vs. delay only, second half | Z = -8.46 | < 0.0001 | **** |
| | | | | Stay vs. delay only, first half | Z = 0.50 | 0.9961 | ns |

| | | | | | | | |
|---|---|---|---|---|---|---|---|
| | | | | Stay vs. delay only, Second half | Z = -0.84 | 0.9594 | ns |
| | | | | Switch vs. stay, first half | Z = -5.43 | <0.0001 | **** |
| | | | | Switch vs. stay, second half | Z = -6.28 | <0.0001 | **** |
| Figure *3*C | Hippocampal position decoding; theta timescale update trial types | Switch: n = 1295 trials; Stay: n = 552 trials; delay only: n = 927 trials | Linear mixed effects model; ANOVA Tukey post-hoc pairwise test | Trial type (switch, delay only), new goal arm | F(5500.03) = 13.96 | < 0.0001 | **** |
| | | | | Switch vs. delay only, first half | Z = -5.26 | < 0.0001 | **** |
| | | | | Switch vs. delay only, second half | Z = -4.94 | < 0.0001 | **** |
| | | | | Stay vs. delay only, first half | Z = -1.36 | 0.7534 | ns |
| | | | | Stay vs. delay only, Second half | Z = -2.76 | 0.0634 | ns |
| | | | | Switch vs. stay, first half | Z = -3.02 | 0.0303 | * |
| | | | | Switch vs. stay, second half | Z = -1.27 | 0.8041 | ns |
| Figure *4*E | Prefrontal choice decoding; posterior probability density integrated by goal arm | Switch: n = 1109 trials, Delay only: n = 798 trials | Linear mixed effects model; ANOVA, Tukey post-hoc pairwise test | Trial type (switch, delay only), choice | F(3777.66) = 78.10 | < 0.0001 | **** |
| | | | | Switch vs. delay only, initial choice | z = 12.58 | < 0.0001 | **** |
| | | | | Switch vs. delay only, new choice | Z = -5.45 | < 0.0001 | **** |
| Figure *4*E | Prefrontal choice decoding; Difference in initial vs. new choice posterior probability density | Switch: n = 1109 trials, Delay only: n = 798 trials | Linear mixed effects model; ANOVA | Trial type (switch, delay only) | F(1896.40) = 147.75 | < 0.0001 | **** |
| Figure *4*F | Hippocampal choice decoding; posterior probability density | Switch: n = 1238 trials Delay only: n = 885 trials | Linear mixed effects model; ANOVA, Tukey post-hoc pairwise test | Trial type (switch, delay only), choice | F(4246.25) = 97.58 | < 0.0001 | **** |
| | | | | Switch vs. delay only, initial choice | Z = 9.17 | < 0.0001 | **** |

| | | | | | | | |
|---|---|---|---|---|---|---|---|
| | integrated by goal arm | | | Switch vs. delay only, new choice | Z = -6.41 | < 0.0001 | **** |
| Figure *4*F | Hippocampal choice decoding; difference in initial vs. new choice posterior probability density | Switch: n = 1109 trials Delay only: n = 798 trials | Linear mixed effects model; ANOVA | Trial type (switch, delay only) | F(2037.93) = 114.34 | < 0.0001 | **** |
| Figure *4*F | Hippocampal choice decoding; difference in initial vs. new | Switch: n = 50 sessions, Delay only: n = 50 sessions | Wilcoxon signed-rank test | Switch vs. zero | W = 577 | 0.5625 | ns |
| | | | | Delay only vs zero | W = 49 | < 0.0001 | ns |
| Figure 5D | Hippocampal position decoding; posterior probability density integrated by goal arm and difference | Switch: n = 1295 trials Stay: n = 551 trials Delay only: n = 926 trials | Linear mixed effects model; ANOVA, Tukey post-hoc pairwise test | Trial type (switch, stay, delay only), goal arm | F(5498.95) = 18.14 | < 0.0001 | **** |
| | | | | Stay vs. switch, initial goal | Z = -5.72 | < 0.0001 | **** |
| | | | | Stay vs. delay only, initial goal | Z = -1.56 | 0.9927 | ns |
| | | | | Stay vs. switch, new goal | Z = -3.05 | 0.0274 | * |
| | | | | Stay vs. delay only, new goal | Z = -1.56 | 0.6267 | ns |
| Figure 5D | Hippocampal position decoding; Difference in initial vs. new goal arm posterior probability density | Switch: n = 1295 trials Stay: n = 551 trials Delay only: n = 926 trials | Linear mixed effects model; ANOVA, Tukey post-hoc pairwise test | Trial type (switch, stay, delay only) | F(2750.69) = 1.69 | 0.1849 | ns |
| | | | | Stay vs. switch | T = -1.79 | 0.1740 | ns |
| | | | | Stay vs. delay only | T = 1.50 | 0.2897 | Ns |
| Figure 5F | Prefrontal choice decoding; posterior probability density integrated by goal arm and difference | Switch: n = 1109 trials, Stay: n = 483 trials, Delay only: n = 798 trials | Linear mixed effects model; ANOVA, Tukey post-hoc pairwise test | Trial type (switch, stay, delay only), choice | F(4739.91) = 55.44 | < 0.0001 | **** |
| | | | | Stay vs. switch, initial choice | Z = 6.77 | < 0.0001 | **** |
| | | | | Stay vs. delay only, initial choice | Z = 3.68 | 0.0032 | ** |
| | | | | Stay vs. switch, new choice | Z = -3.46 | 0.0072 | ** |

| | | | | | | | |
|---|---|---|---|---|---|---|---|
| | | | | Stay vs. delay only, new choice | Z = -1.29 | 0.7914 | ns |
| Figure 5F | Prefrontal choice decoding; Difference in initial vs. new choice posterior probability density | Switch: n = 1109 trials, Stay: n = 483 trials, Delay only: n = 798 trials | Linear mixed effects model; ANOVA, Tukey post-hoc pairwise test | Trial type (switch, stay, delay only) | F(2370.62) = 76.21 | < 0.0001 | **** |
| | | | | Stay vs. switch | T = 6.86 | < 0.0001 | **** |
| | | | | Stay vs. delay only | T = 3.25 | 0.0033 | ** |
| Figure 6A | Hippocampal position decoding; posterior probability density integrated by goal arm | Switch, correct: n = 1295 trials, Switch, incorrect: n = 407 trials | Linear mixed effects model; ANOVA, Tukey post-hoc pairwise test | Trial outcome (correct, incorrect), goal arm | F(3283.78) = 4.99 | 0.0019 | ** |
| | | | | Correct vs. incorrect, initial goal | Z = 1.75 | 0.2967 | ns |
| | | | | Correct vs. incorrect, New goal | Z = 3.51 | 0.0025 | ** |
| Figure 6A | Hippocampal position decoding; difference in initial vs. new goal arm posterior probability density | Switch, correct: n = 1295 trials, Switch, incorrect: n = 407 trials | Linear mixed effects model; Tukey post-hoc pairwise test | Correct vs. incorrect, initial vs. new | T(1602.30) = -0.89 | 0.3725 | ns |
| Figure 6A | Hippocampal position decoding; pre update | Switch, correct: N = 1292 trials, switch, incorrect: n = 407 trials | Linear mixed effects model; ANOVA, Tukey post-hoc pairwise test | Trial outcome (correct, incorrect), goal arm | F(3162.37) = 7.62 | < 0.0001 | **** |
| | | | | Correct vs. incorrect, initial goal | Z = -4.17 | 0.0002 | *** |
| | | | | Correct vs. incorrect, new goal | Z = 2.27 | 0.1058 | ns |
| Figure 6B | Prefrontal choice decoding; posterior probability density | Switch, correct: N = 1109 trials, switch, incorrect: n = 359 trials | Linear mixed effects model; ANOVA, Tukey post-hoc pairwise test | Trial outcome (correct, incorrect), choice | F(2754.10) = 8.29 | < 0.0001 | **** |
| | | | | Correct vs. incorrect, initial choice | T = -1.26 | 0.5874 | ns |

| | | | | | | | |
|---|---|---|---|---|---|---|---|
| | integrated by goal arm | | | Correct vs. incorrect, New choice | T = 4.14 | 0.0002 | *** |
| Figure 6B | Prefrontal choice decoding; difference in initial vs. new goal arm posterior probability density | Switch, correct: N = 1109 trials, switch, incorrect: n = 359 trials | Linear mixed effects model; Tukey post-hoc pairwise test | Correct vs. incorrect, initial vs. new choice | T = -3.91 | 0.0001 | **** |
| Figure 6B | Prefrontal choice decoding; pre update | Switch, correct: N = 1107 trials, switch, incorrect: n = 359 trials | Linear mixed effects model; ANOVA, Tukey post-hoc pairwise test | Trial outcome (correct, incorrect), choice | F(2808.03) = 7.98 | < 0.0001 | **** |
| | | | | Correct vs. incorrect, initial choice | t = -2.54 | 0.0536 | ns |
| | | | | Correct vs. incorrect, new choice | T = 0.34 | 0.9862 | ns |
| Figure 6C | Hippocampal position prediction accuracy | Switch: n = 1702 trials, n = 1000 shuffles | Permutation test | Switch post update vs. shuffled trials | N/A | < 0.0001 | **** |
| Figure 6C | Prefrontal choice prediction accuracy | Switch: n = 1468 trials, n = 1000 shuffles | Permutation test | Switch post update vs. shuffled trials | N/A | < 0.0001 | **** |
| Figure 7B | Hippocampal position vs. view angle at update onset | N = 1235 trials | Spearman correlation | View angle vs. initial goal codes | Rho = 0.1540 | < 0.0001 | **** |
| | | N = 1235 trials | Spearman correlation | View angle vs. new goal codes | Rho = 0.0261 | 0.3603 | ns |
| Figure 7C Figure 7 | Prefrontal choice vs. view angle at update onset | N = 1103 trials | Spearman correlation | View angle vs. initial choice codes | Rho = 0.3344 | < 0.0001 | **** |
| | | N = 1103 trials | Spearman correlation | View angle vs. new choice codes | Rho = -0.1814 | < 0.0001 | **** |

***Table 2*** **Statistical analysis details.**

Details for all statistical comparisons. Statistical significance abbreviations: ns (not significant) $P > 0.05$, *$P < 0.05$, **$P < 0.01$, ***$P < 0.001$, ****$P<0.0001$. Linear mixed effects models were generated using nested random effects for animal and recording session levels. When comparing trial types, linear mixed effects models included delay, update, and stay trials to compare each trial type with the same model.

### SUPPLEMENTARY FIGURES

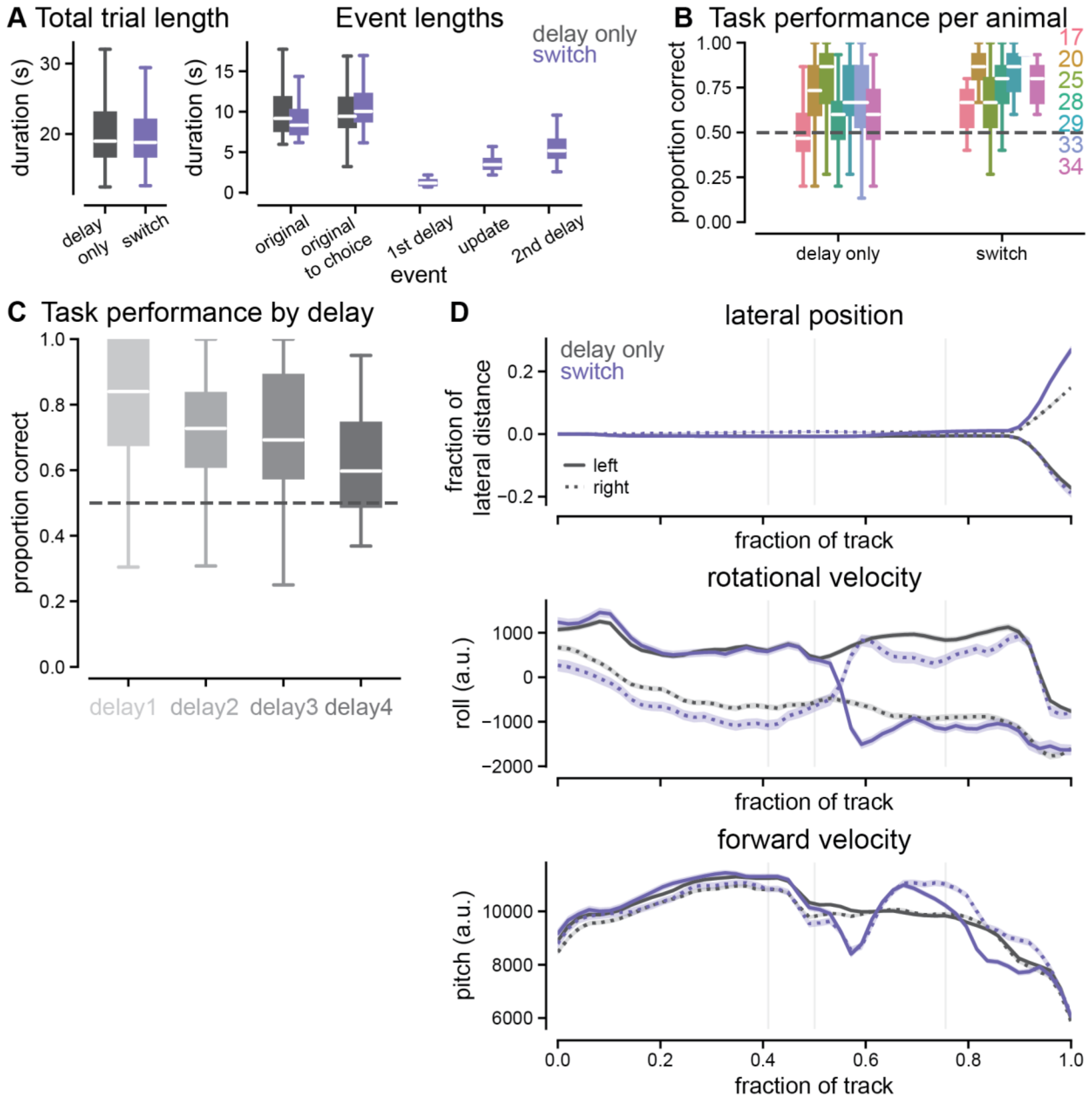


***Supplementary Figure 1.*** **Additional update task behavioral metrics and task details.**

**A.** Durations of events during behavioral task across trial types. *Left*, total trial duration from trial start to choice being made (Delay only: black, switch: purple). (delay only: 20.93 ± 0.09 seconds, n = 7313 trials, percentiles = 12.48, 16.83, 18.92, 22.89, 259.97, switch: 20.79 ± 0.23 seconds, n = 1886 trials, percentiles = 12.66, 16.81, 18.76, 21.86, 240.23). Right, durations for different cue phases of the task across trial types. Original cue to choice indicates period from original cue offset until choice made, equivalent to the duration of the delay phase for delay only trials. No values recorded for 1st delay, update, and 2nd delay for delay only trials since their were no cue updates and the duration of the delay is captured by the 'original to choice made' metric.

**B.** Performance across trial types by individual animal during recording sessions. Colored numbers indicate animal identification number in color corresponding to bar graph. See Table 1 for individual animal metadata.

**C.** Performance for delay only trials across different delay durations during the warmup training at the start of each session.

**D.** Trajectories of additional behavioral metrics across delay only (black) and switch(purple) trials. Note that the lateral position is restricted to a very small distance for the center arm of the environment.

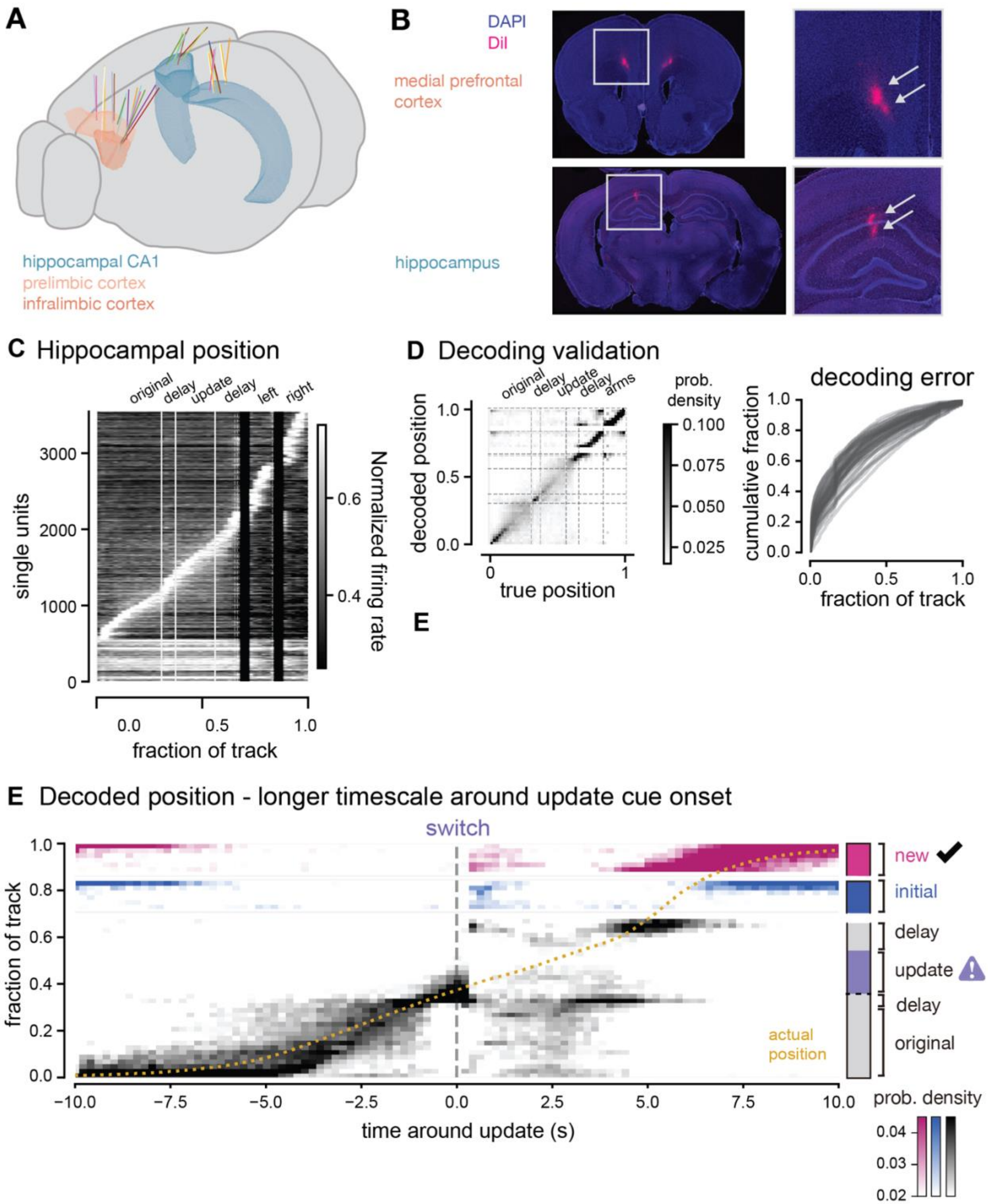


***Supplementary Figure 2.*** **Recording location details and bayesian decoding validation for hippocampal position**

**A.** Reconstructed electrode insertion paths from histological data registered to Allen mouse atlas. Each line indicates a separate electrode shank or probe if individual shanks could not be isolated. Hippocampal CA1, prelimbic cortex, and infralimbic cortex regions highlighted.

**B.** Example histological images showing electrode locations (indicated by DiI that was coated on probe) in brain with cells stained with DAPI to visualize structures.

**C.** Tuning curves for position of all units from hippocampal CA1 built from encoding model for Bayesian decoder (80% of the delay only trials). Units are sorted by place field peak location, bottom units have no significant place fields.

**D.** *Left,* Confusion matrices (decoded position from hippocampal neural activity on y-axis compared to actual position from behavioral data on x-axis) for delay only trials. Dashed lines indicate locations in the environment. *Right*, Decoding error distributions for each recording session, error calculated as absolute difference between true and peak decoded position.

**E.** *Left,* Decoding output (posterior probability density) before and after the update cue is presented at a longer timescale (10 seconds before and after). The heatmap shows transient increases in non-local (pink and blue) and decreases in local (grey) representations after the update cue. Average across all recording sessions for switch trials. Heatmap indicates which positions are most strongly decoded by the spiking activity of all hippocampal neurons. Pink and blue indicate decoding of positions within the new and initial goal arms, respectively (including the entire goal arms after the choice point). Animals' actual position (yellow dashed line) shown over the same time window. Data from both left and right trial types were combined. Two-dimensional position in the virtual track was converted to a 1D position (see ***Methods***) to disambiguate the new and initial arms of the environment. *Right*, schematic of locations of the task as shown on y-axis of left panels.

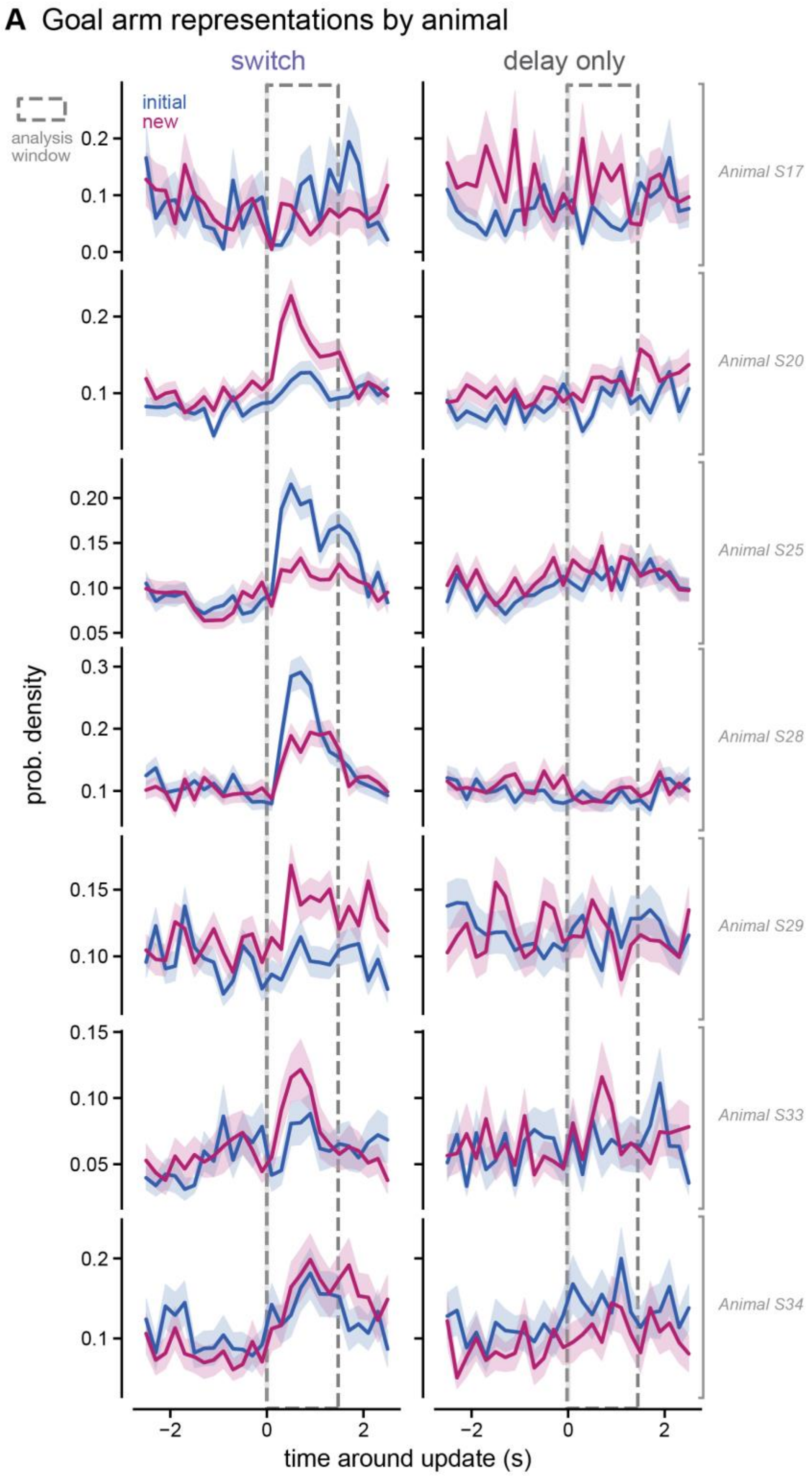


***Supplementary Figure 3. Hippocampal position decoding in each animal.***

***A.*** Integrated probability densities of the new (pink) and initial (blue) goal arms around the update cue on switch (left) and delay only (right) trials for each animal (row). Mean ± SEM across all trials shown. Dashed window indicates analysis window used for statistical quantification.

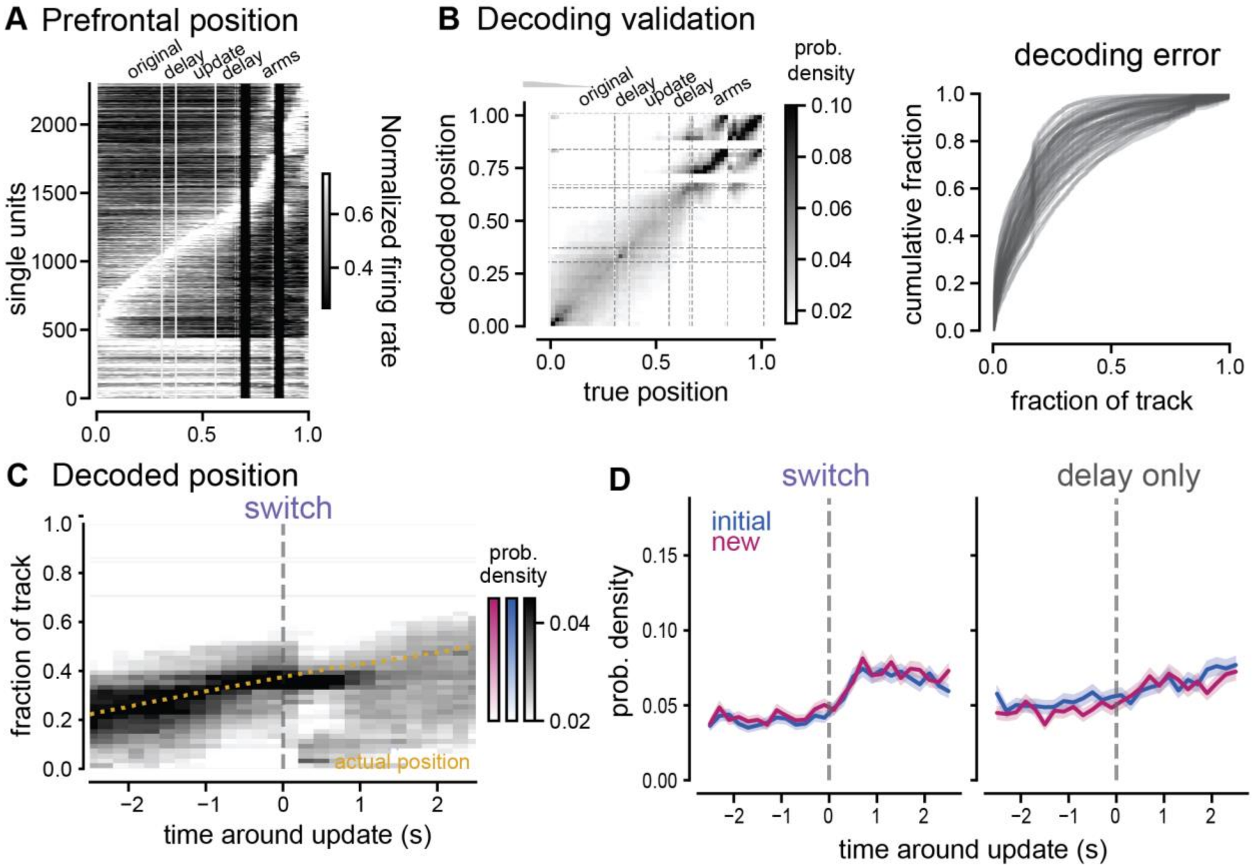


***Supplementary Figure 4.*** **Bayesian decoding of position for prefrontal cortex**

- **A.** Tuning curves for position of all units from medial prefrontal cortex built from encoding model for Bayesian decoder (80% of the delay only trials).
- **B.** *Left,* Confusion matrix (decoded position from prefrontal neural activity on y-axis compared to actual position from behavioral data on x-axis) for delay only trials. *Right,* cumulative distribution of decoding errors where error is measured as the absolute difference between the actual position and the peak decoding position.
- **C.** Decoding output (probability density) from prefrontal cortex before and after the update cue is presented on switch trials, average for all recording sessions. Heatmap indicates stronger likelihood of those position being decoded by the spiking activity of all prefrontal cortex neurons. Animals' actual position (yellow dashed line) shown over the same time window.
- **D.** Integrated probability densities of the new (pink) and initial (blue) goal arms around the update cue on switch and delay only trials.

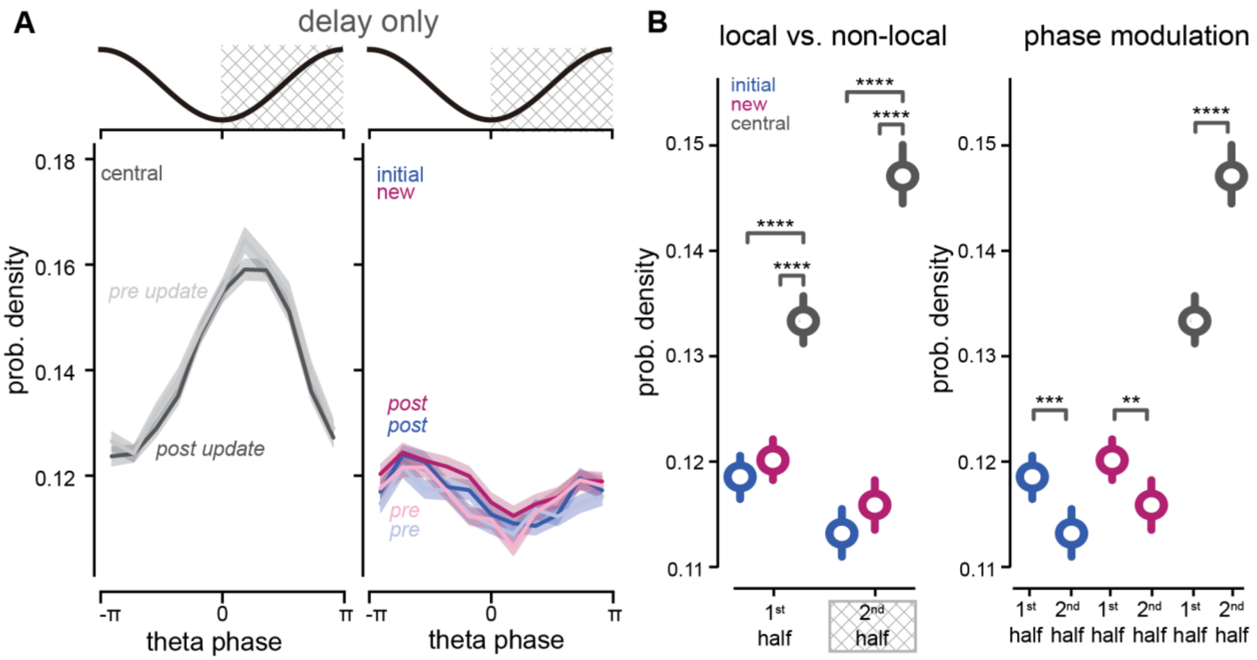


***Supplementary Figure 5.*** **Theta phase modulation of position representations for stay and delay only trials**

A. *Top,* theta phase with cross hatching indicating the second half of the theta cycle. *Bottom, Left,* average decoded central arm (black and grey) posterior probability densities by theta phase pre (-1.5 to 0 seconds, lighter shades) and post (0 to 1.5 seconds, darker shades) the update cue onset for delay only trials. *Right*, as in left for new (pink) and initial (blue) goal arms. Mean ± SEM across all trials shown.

B. *Left,* probability density of the central (local), new (non-local), and initial (non-local) arm in the first and second halves ($1^{st}$ half indicates $-\pi$ to 0 and $2^{nd}$ half indicates 0 to $\pi$) of theta cycle after the update cue onset on switch trials. *Right,* as on left with statistical comparisons between phases. Initial, $-\pi$ to 0 : 0.12 ± 0.00, n = 927 trials, percentiles = 0.00, 0.10, 0.12, 0.14, 0.42; initial, 0 to $\pi$: 0.11 ± 0.00, n = 927 trials, percentiles = 0.00, 0.09, 0.11, 0.13, 0.35; new, $-\pi$ to 0: 0.12 ± 0.00, n = 927 trials, percentiles = 0.00, 0.10, 0.12, 0.14, 0.32; new, 0 to $\pi$: 0.12 ± 0.00, n = 927 trials, percentiles = 0.00, 0.09, 0.11, 0.13, 0.38; central, $-\pi$ to 0: 0.13 ± 0.00, n = 927 trials, percentiles = 0.00, 0.11, 0.13, 0.15, 0.34; central, 0 to $\pi$: 0.15 ± 0.00, n = 927 trials, percentiles = 0.00, 0.12, 0.14, 0.16, 0.39. Linear mixed effects model (LME) and ANOVA Tukey post-hoc, see Table 2 for statistical details.

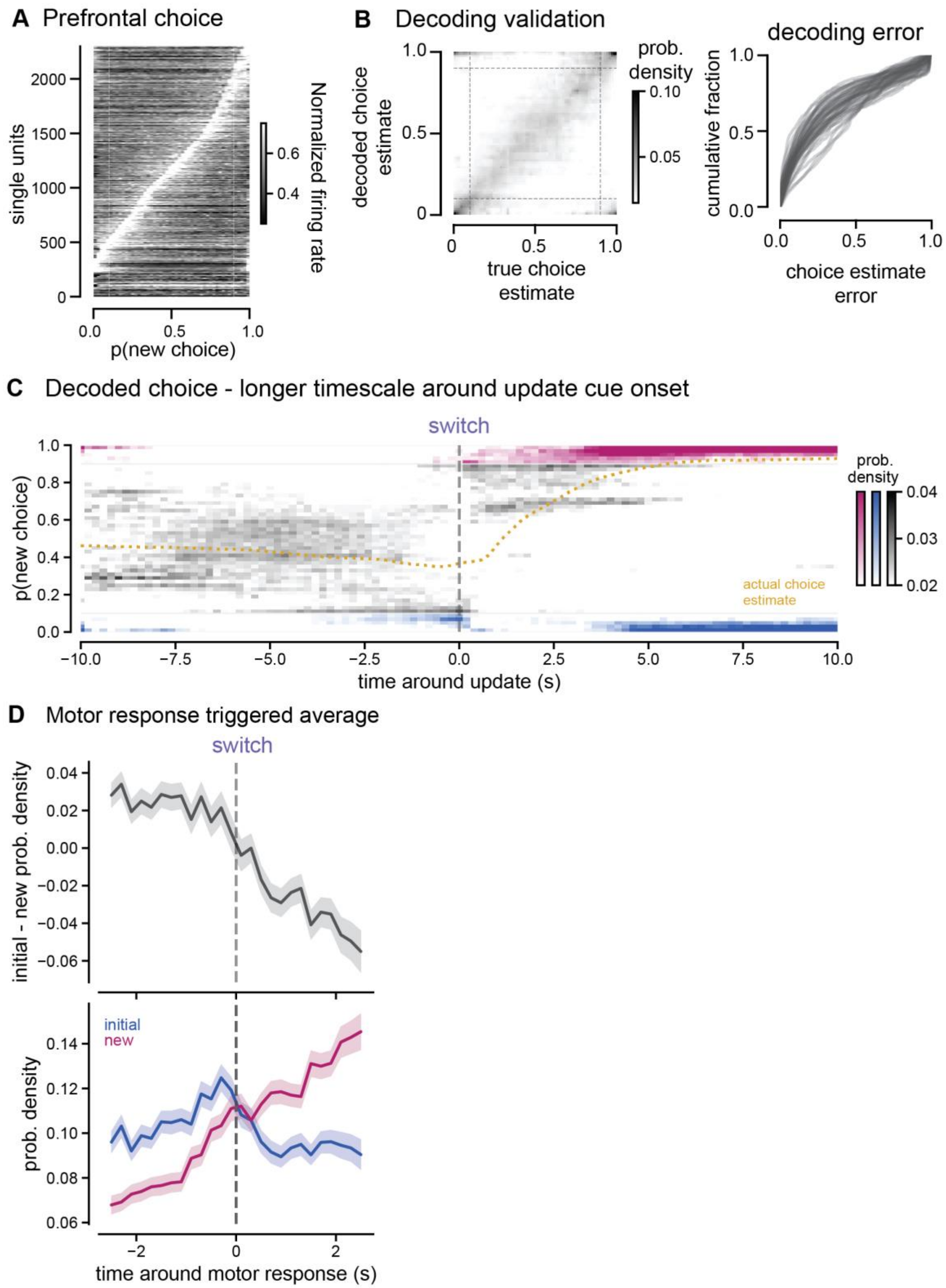


***Supplementary Figure 6.*** **Bayesian decoding validation of prefrontal choice representation.**

- **A.** Tuning curves for choice of all units from medial prefrontal cortex built from encoding model for Bayesian decoder (80% of the delay only trials). Units sorted by firing rate peak.

**B.** *Left,* Confusion matrix (decoded choice from prefrontal neural activity compared to actual choice estimate from behavioral data) for all delay only trials. *Right,* cumulative distribution of decoding errors where error is measured as the absolute difference between the actual position and the peak decoding position.

**C.** Decoding output (posterior probability density) before and after the update cue is presented at a longer timescale (10 seconds before and after). The heatmap shows a rapid decrease in the initial choice (blue) and increase in the new choice (pink) represnetations after the update cue. Average across all recording sessions for switch trials. Heatmap indicates which choices are most strongly decoded by the spiking activity of all prefrontal neurons. Blue and pink indicate decoding of initial and new choices, respectively. Animals' actual choice estimate (yellow dashed line) shown over the same time window. Data from both left and right trial types were combined.

**D.** Neural activity aligned to motor response times on an individual trial basis. Motor responses were identified as the moment the rotational velocity began decelerating (the differential of the velocity signal switched from positive to negative). *Top*, difference between initial and new choice decoding outputs. *Bottom*, initial and new choice decoding outputs.

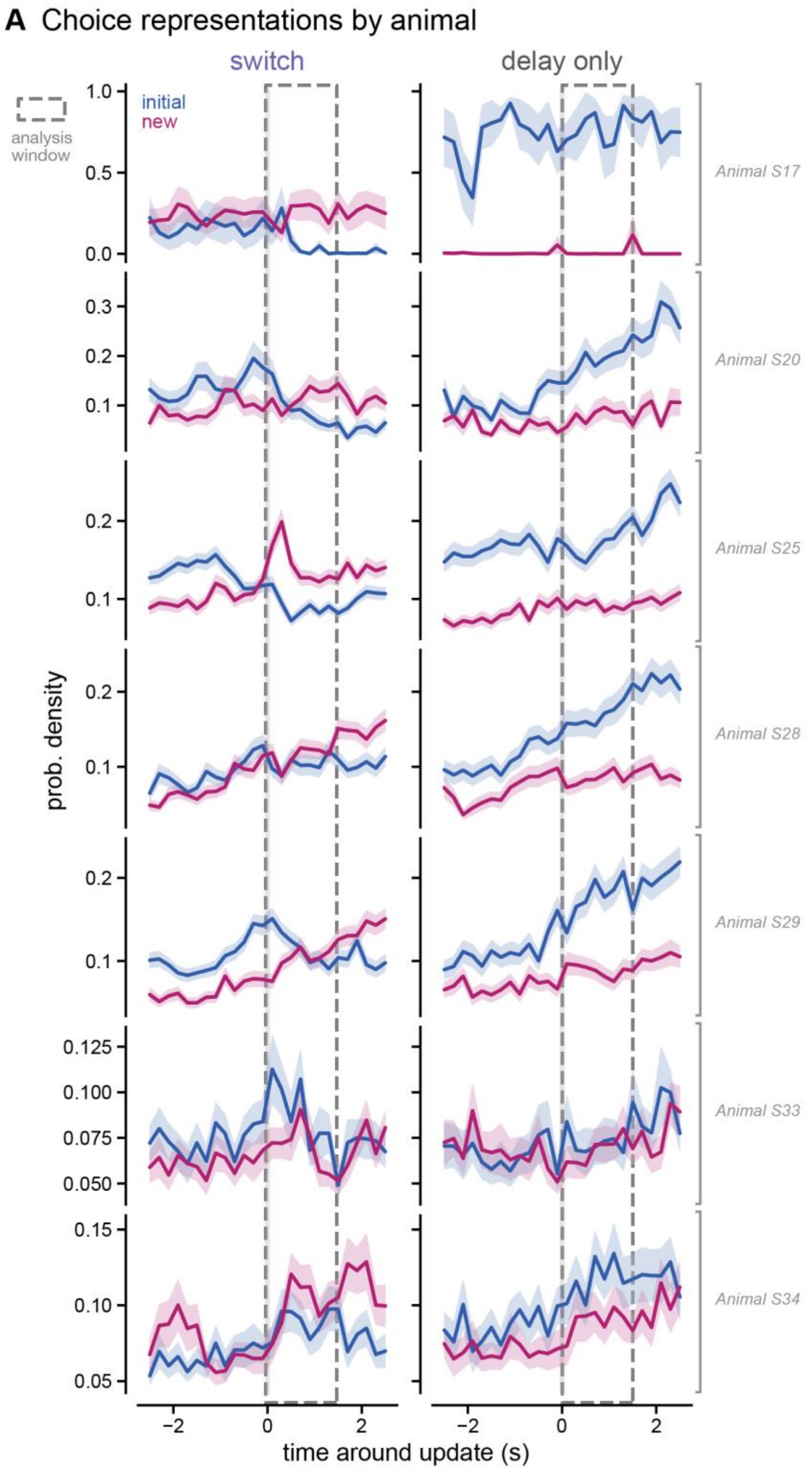


***Supplementary Figure 7. Prefrontal choice decoding in each animal.***

**A.** Integrated probability densities of the new (pink) and initial (blue) choice around the update cue on switch (left) and delay only (right) trials for each animal (row). Mean ± SEM across all trials. Dashed window indicates analysis window used for statistical quantification.

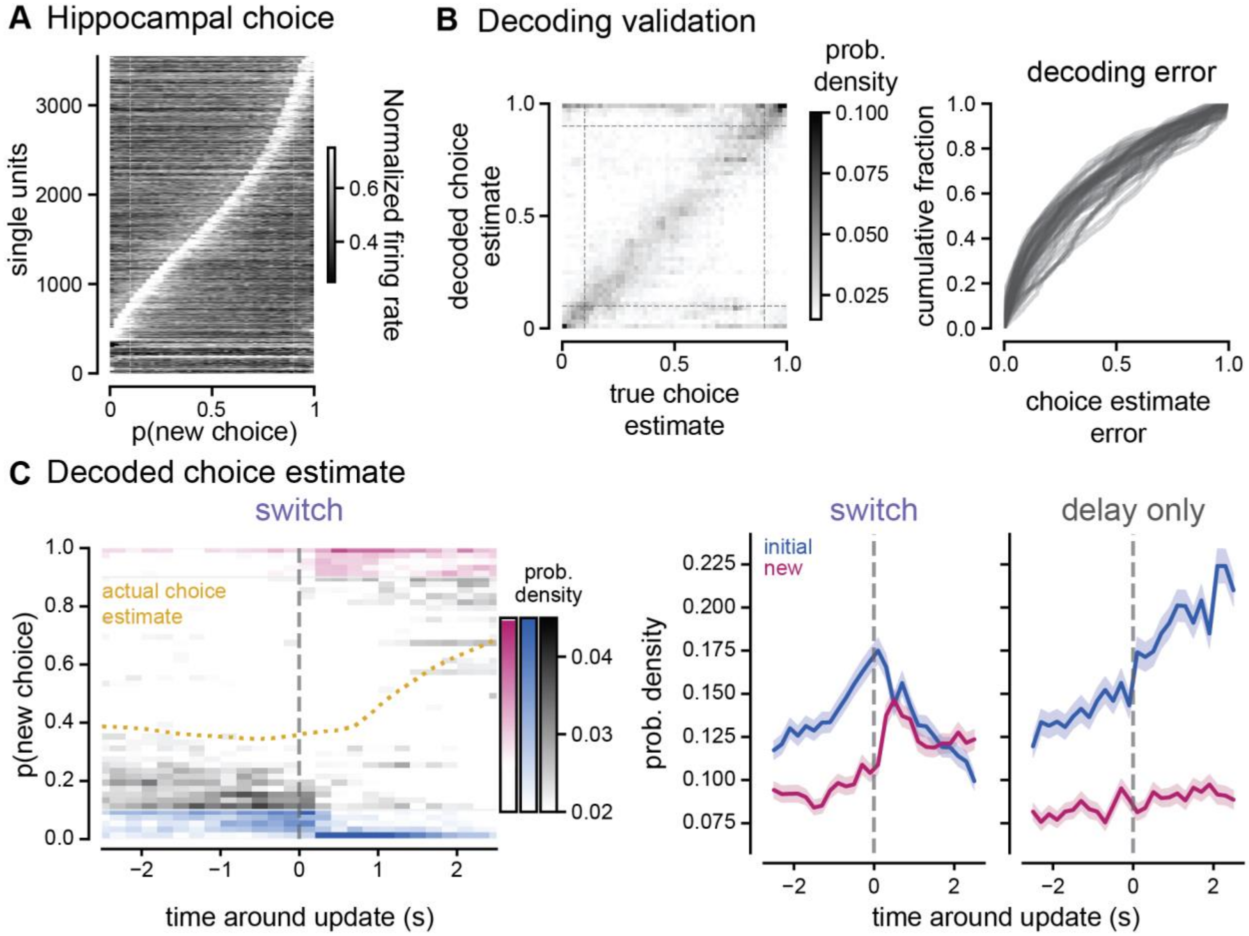


***Supplementary Figure 8.*** **Bayesian decoding of choice representation in hippocampus.**

- **A.** Tuning curves for choice of all units from hippocampus. Tuning curves built from encoding model for Bayesian decoder (80% of the delay only trials).
- **B.** *Left,* Confusion matrix (decoded choice from hippocampal neural activity compared to actual choice estimate from behavioral data) for delay only trials. *Right,* cumulative distribution of decoding errors where error is measured as the absolute difference between the actual position and the peak decoding position.
- **C.** *Left,* decoding output (probability density) from hippocampus before and after the update cue is presented on switch trials, average for all recording sessions. Heatmap indicates stronger likelihood of those choices being decoded by the spiking activity of all hippocampal neurons. Animals actual choice estimate (yellow dashed line) shown over the same time window. *Right,* Integrated probability densities of the new (pink) and initial (blue) choices around the update cue on switch trials. Mean ± SEM shown.

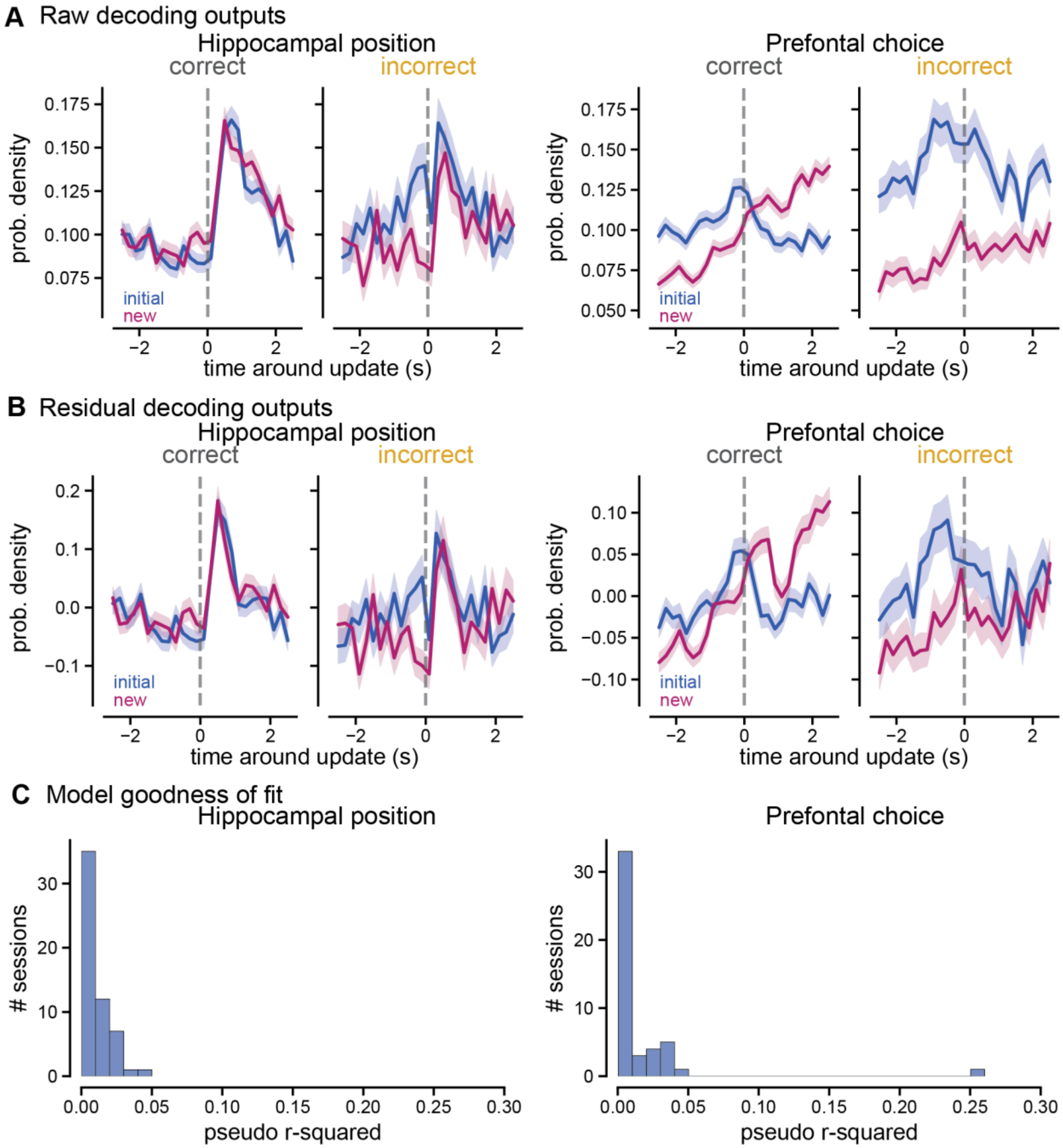


***Supplementary Figure 9.*** **Generalized linear model inputs and outputs to control for differences in velocity.**

**A.** *Left,* raw integrated probability densities of the new (pink) and initial (blue) position estimates around the update cue on correct (grey) and incorrect (yellow) trials from hippocampal CA1. *Right*, raw integrated probability densities of the new (pink) and initial (blue) choice estimates around the update cue on correct (grey) and incorrect (yellow) trials from prefrontal cortex. Mean ± SEM across all trials shown.

**B.** To account for the effects of speed and direction on the differences in neural codes between correct and incorrect trials, we applied a GLM with rotational and translational velocities as the explanatory variables or predictors and the goal decoding output as the response variable. As in **A** for residuals of GLM.

**C.** Model goodness of fit. Distribution of pseudo r-squared values (one per session) as an estimate of how much the velocity explains the neural outputs.

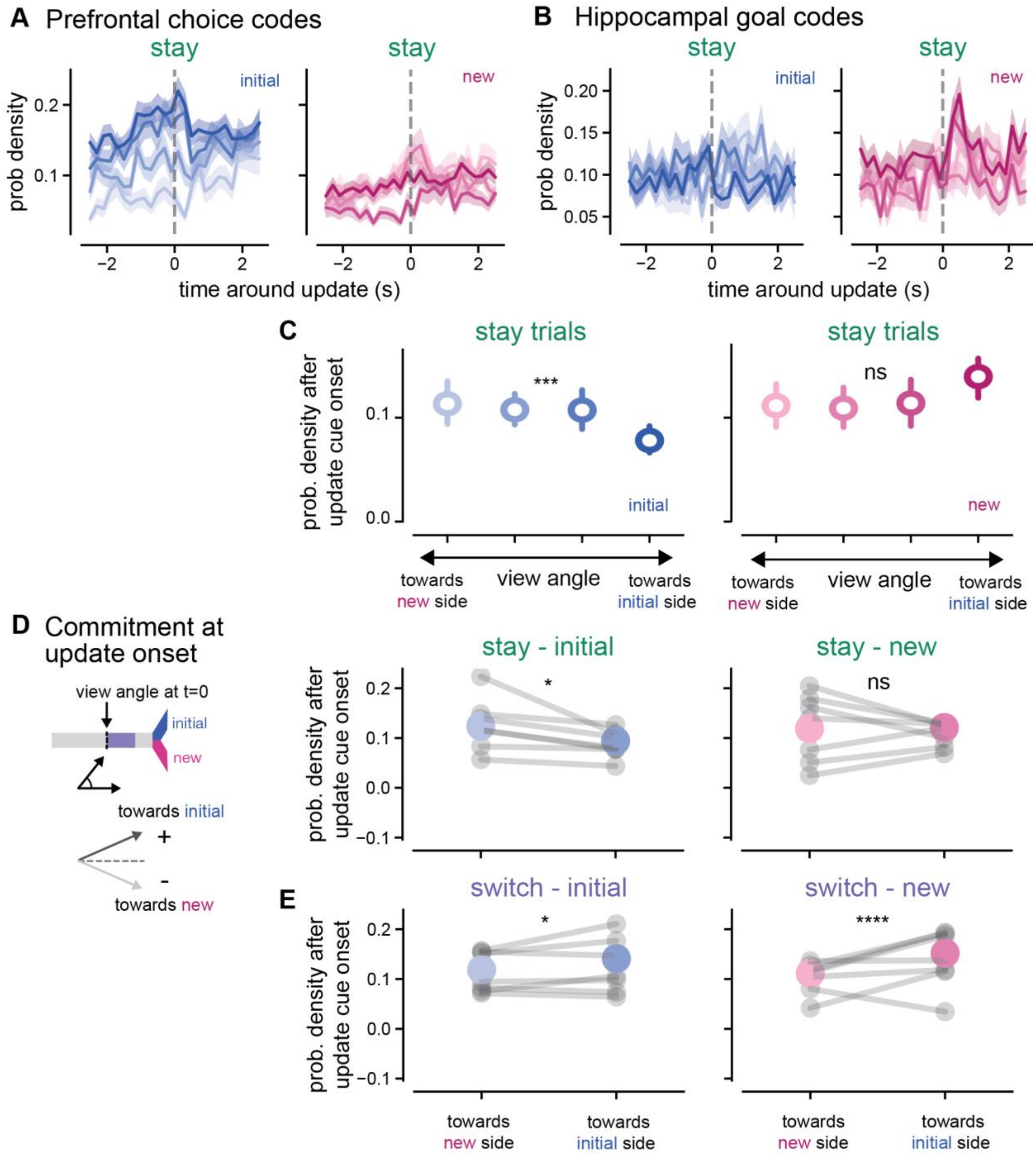


***Supplementary Figure 10*. Choice commitment breakdown of decoding outputs across trial types**

**A.** Prefrontal cortex choice codes with trials binned by commitment quartiles based on view angle at the time of update cue onset for stay trials. *Left*, initial goal representation (blue), *Right*, new goal representation (pink). Darker colors indicates trials with a stronger commitment to the initial side measured by a greater view angle towards the initial side at update cue onset, lighter colors indicates weaker commitment to the initial side, measured by a view angle towards the new side at update cue onset.

**B.** As in **A** for hippocampal goal location codes on stay trials

**C.** Average probability density after the update across all view angle quartiles and separated by higher view angle values indicating strong commitment to the initial side (darker shades) to lower view angle values indicating weaker commitment to the initial side (lighter shades). *Left*, initial goal representations in hippocampus in blue. *Right,* new goal representations in hippocampus in pink.

**D.** *Left,* schematic of commitment estimate breakdown from view angle at the onset of the update cue. *Right,* average probability density separated into all trials with view angles towards the initial side vs. all trials with view angles towards the new side at the time of the update cue on stay trials. Initial goal representations in left column in blue, new goal representations in right column in pink (initial goal representations, towards new side: 0.12 ± 0.01, n = 83 trials, percentiles = 0.00, 0.04, 0.09, 0.16, 0.58; initial goal representations, towards initial side: 0.09 ± 0.00, n = 443 trials, percentiles = 0.00, 0.01, 0.06, 0.14, 0.49; new goal representations, towards new side: 0.12 ± 0.01, n = 83 trials, percentiles = 0.00, 0.03, 0.09, 0.18, 0.44; new goal representations, towards initial side: 0.12 ± 0.01, n = 443 trials, percentiles = 0.00, 0.02, 0.09, 0.18, 0.67).

**E.** As in **D** for switch trials (initial goal representations, towards new side: 0.12 ± 0.01, n = 362 trials, percentiles = 0.00, 0.02, 0.09, 0.18, 0.87; initial goal representations, towards initial side: 0.14 ± 0.00, n = 873 trials, percentiles = 0.00, 0.03, 0.10, 0.20, 0.74; new goal representations, towards new side: 0.11 ± 0.01, n = 362 trials, percentiles = 0.00, 0.02, 0.07, 0.16, 0.68; new goal representations, towards initial side: positive: 0.15 ± 0.00, n = 873 trials, percentiles = 0.00, 0.04, 0.11, 0.22, 0.97).
